# Testis-specific H2BFWT disrupts nucleosome integrity through reductions of DNA-histone interactions

**DOI:** 10.1101/2022.07.20.500751

**Authors:** Dongbo Ding, Matthew Y.H. Pang, Mingxi Deng, Thi Thuy Nguyen, Xulun Sun, Zhichun Xu, Yingyi Zhang, Yuanliang Zhai, Yan Yan, Toyotaka Ishibashi

**Affiliations:** Division of Life Science, The Hong Kong University of Science and Technology, Clear Water Bay, NT, HKSAR, China; School of Biological Sciences, The University of Hong Kong, Pokfulam, Hong Kong, HKSAR, China

**Keywords:** histone variant, spermatogenesis, testis-specific, optical tweezers, H2BFWT

## Abstract

During spermatogenesis, multiple testis-specific histone variants are involved in the dynamic chromatin transitions. H2BFWT is a primate testis-specific H2B variant with hitherto unclear functions, and SNPs of H2BFWT are closely associated with male non-obstructive infertility. Here, we found that H2BFWT is preferentially localized in the sub-telomeric regions and the promoters of genes highly expressed in testis from differentiated spermatogonia to early spermatocytes. Cryo-EM structural analysis shows that H2BFWT nucleosomes are defined by weakened interactions between H2A-H2BFWT dimer and H4, and between histone octamer and DNA. Furthermore, one of its SNPs, H2BFWTH100R further destabilizes nucleosomes and increases the nucleosome unwrapping rate by interfering with the interaction with H4K91. Our results suggest that H2BFWT may be necessary for the regulation of spermatogenesis-related gene expression by decreasing transcriptional barriers, and that H2BFWTH100R overdrives its nucleosome-destabilizing effects which causes infertility.

## Introduction

A nucleosome consists of 147 bp DNA wrapping around a histone octamer, which contains two copies of histone H2A, H2B, H3, and H4. Linker histone H1/H5 binds to adjacent nucleosomes and helps to form chromatin fiber, which is one of the key factors for gene expression regulation ^1,2^. In addition to canonical core histone H2A, H2B, H3, and H4, there are multiple histone variants, such as H2A.Z, H2A.X, H2A.B (H2A.Bbd), macroH2A, TH2A, TH2B, H2BFWT, CENPA, H3.3, H3T (H3.4), H3.5, H3.X, H3.Y and H4G expressed in human ^3–5^. These histone variants possess distinct functional features from their canonical counterparts highlighted by their unique localization, structure, and/or expression pattern ^6^.

Spermatogenesis is the process in which diploid germ stem cells develop into haploid sperms via meiosis. The chromatin structures undergo dynamic changes during spermatogenesis. Specifically, canonical histones and somatic histone variants are replaced by testis-specific histone variants, such as H3T, H3.5, H2A.B, TH2A, TH2B, and H2BFWT, in a tightly controlled process ^7,8^. For example, H3T is one of the testis-specific H3 variants and has only four amino acid residues different from its canonical counterpart H3.1 ^9^. Largely due to the two substitutions in positions M71 and V111 of H3.1, H3T-containing nucleosomes have much lower stability than canonical nucleosomes ^9^. Furthermore, the V111 substitution in H3T weakens its interaction with histone chaperone Nap1L1, while allowing H3T to bind strongly to Nap1L2 ^10^. The mouse homolog of H3T, H3t, which does not contain M71 and V111, replaces the canonical H3 histones during differentiating spermatogonia in mice ^11^. The *H3t* knockout mouse had smaller testis and exhibited azoospermia due to loss of haploid germ cells ^11^. H3.5, another testis-specific histone H3 variant only present in hominids, is expressed around meiosis I, but not in mature sperm ^12,13^. The H3.5 amino acid sequence has only five amino acid residue alterations from H3.3. Compared with the H3.3 nucleosome, the H3.5 nucleosome is less stable, and the L104 site of H3.5 contributes most to this decreased nucleosome stability ^12,13^. ChIP-sequencing results of human testicular cells demonstrated an enrichment of H3.3 in the transcription start site (TSS) of active genes, while H3.5 accumulates around the TSS of both active and inactive genes ^12^.

Mammalian testis-specific histone H2A variant, H2A.B is known as one of the fast-evolving histones and is encoded by a gene in the X chromosome ^14,15^. Nucleosomes containing H2A.B are destabilized and only protect 118±2 bp of DNA ^16^. H2A.B is expressed in the later stages of round spermatid, with enrichment at the TSS and the beginning of the gene body of actively transcribed genes ^17,18^. In addition to its involvement in pre-mRNA processing, H2A.B enables activation of previously repressed genes ^17–19^ by associating with active RNA polymerase II (Pol II) and transcription elongation factors ^18–20^.

TH2B and H2BFWT are testis-specific histone H2B variants ^7,8^. Compared with canonical nucleosomes, nucleosomes containing both TH2A and TH2B have weaker DNA-histone interactions, and there are also some changes in the intra-nucleosome TH2A-TH2A interaction at the TH2A L1 loop region. Because of these changes, the TH2A/TH2B nucleosome has lower stability than the canonical nucleosome ^21^. The expression of TH2B starts in early spermatocytes and lasts until round spermatids in mice, and it replaces H2B completely by the secondary spermatocyte stage ^22^. ChIP-sequencing analysis showed that TH2B is involved in the process of chromatin transition from nucleosome to protamine ^22^. TH2B was found to be depleted from the TSS regions, and its genomic localization was negatively correlated with H2A.Z ^22^. When the *TH2A* and *TH2B* genes were disrupted in mice, mice spermatogenesis was defective ^23^.

H2BFWT (H2BW, H2B histone family member W, testis-specific) is another H2B variant specifically expressed in testis ^24^. Unlike other testis-specific histone variants, H2BFWT is only present in primates ^24^. The *H2BFWT* gene is located at Xq22.2 and contains two introns in humans ^24^. There are two potential AUG start codons in H2BFWT mRNA, however, only the shorter version (152 amino acid residues excluding the methionine) was detected *in vivo* ^25^. H2BFWT represents the shorter 152-amino-acid version in this manuscript if not indicated otherwise. The GFP-conjugated H2BFWT175 (longer version, 175 amino acid residues including the methionine) partially colocalized with telomeric DNA regions in V79 cell ^24^. Compared with the canonical nucleosome, the H2BFWT nucleosome was more sensitive to the SWI/SNF remodeling complex and thus these results could reflect alternation of the histone-DNA interaction ^25^. The highly divergent N-terminal tail of H2BFWT is unable to recruit chromosome condensation factors and therefore, H2BFWT cannot participate in the assembly of mitotic chromosomes ^25^. It was reported that the single-nucleotide polymorphisms (SNPs) in the 5’UTR (−9C>T, which will not produce H2BFWT protein due to creating the ATG codon at −10 and causing a frameshift termination) and gene body (368A>G, which will change the amino acid from H to R at position 100.) are associated with spermatogenesis impairment in different demographic groups ^26–28^. However, H2BFWT functions in spermatogenesis and its mechanistic relationship with infertility remains unknown.

In this study, we found that H2BFWT is mainly present in differentiated spermatogonia and early spermatocytes. H2BFWT is localized at the sub-telomeric regions and enriched near a number of genes highly expressed in testis. We determined the cryo-EM structures of the H2BFWT nucleosomes. From the structure, we found that H2BFWT destabilizes nucleosomes by reducing interactions between H2A-H2BFWT dimer and H4, H2A-H2BFWT dimers and DNA, as well as interactions between H3-H4 tetramer and DNA. Moreover, the shape of the H2A L1 loop is changed and a new hydrogens-bond (H-bond) is formed between the N38 residues (H2A-N38 and H2A’-N38) of the two H2A copies. In addition, we performed single-molecule optical tweezers experiments and found that H2BFWT nucleosomes exhibit a diminished rewrapping rate and an increased unwrapping rate, which translate to an approximately ~40% destabilization in the H2A/H2B-DNA interaction region. Furthermore, H2BFWTH100R, which is more frequently observed in infertile male patients, decreases the stability of the nucleosome further by changing the surface electrostatistic property to positive charge which repulses the nearby positively charged H4K91. Finally, our *in vitro* Pol II transcription assay showed that Pol II can transcribe through H2BFWT nucleosomes more efficiently. Altogether, our data unravels the mechanisms of how H2BFWT functions in the process of early spermatogenesis to regulate spermatogenesis-related gene expression through opening of nucleosome structure.

## Material and Methods

### Human testis immunohistochemistry and immunofluorescence staining

Human Formalin-Fixed, Paraffin-Embedded (FFPE) normal testis tissue slides were obtained from Zyagen (United States) from a 29-year-old donor. To remove all paraffin, testis sections were first washed with xylene and then rinsed with an ethanol/water series for rehydration. Epitope retrieval was achieved by keeping the slides submerged at 98°C for 15 min in sodium citrate buffer (10 mM sodium citrate (pH6.0), 0.05% Tween-20). The slides were then rinsed by water and PBS. After blocking with rabbit serum, the slides were washed with PBS and incubated with rabbit anti-H2BFWT antibody (ab185682, 1/100 dilution for IHC and 1/25 dilution for IF) for 1 h at room temperature. For TH2B staining, a rabbit anti-hsTH2B (1:1500 dilution) antibody was made based on a peptide epitope (amino acid residues 2-17, Figure S1A, Shanghai Youke Biotechnology), and the anti-H4 antibody (ab7311, 1/1000 dilution) was used as the overall positive control. For immunohistochemistry staining, the staining results were visualized using a rabbit specific HRP/DAB (ABC) detection IHC kit (ab64261) according to the manufacturer’s protocol. For immunofluorescence staining, the slides were further incubated with Alexa Fluor™ 568 goat-anti-rabbit secondary antibody (1/800 dilution, A-11036) for 1 h at room temperature. DNA was then stained with Hoechst 33342 and the images were captured using a LSM 980 (ZEISS) confocal microscope.

### Chromatin immunoprecipitation from human testis, sequencing library preparation and sequencing analysis

The ChIP assay from human testis was followed by the published methods with modification ^29,30^. The detailed method can be found in the supplemental information. Briefly, human testis FFPE tissue (Origene, CB811079) was first sliced, then paraffin was removed. After the treatment of MNase to digest the DNA, Thermolabile Proteinase K (NEB, P8111S) with 15 mM EDTA was added to the reaction. The samples were centrifuged and the supernatant was diluted by the ChIP buffer (30 mM Tris-HCl (pH 7.5), 50 mM NaCl, 3 mM EDTA, 0.1 mM PMSF). The diluted sample was mixed with H2BFWT antibody and Protein A/G Plus agarose, and the DNA was extracted and then DNA libraries were prepared with Smarter Thruplex DNAseq Kit (Clontech, R400674) following the manufacturer’s protocols with amplification for 10-13 circles. Sequencing was performed based on Nextseq 500 instrument (Illumina) with paired-end 150 bp (PE150) sequencing method. The sequence data was analyzed by standard analysis method (detail method can be found in the supplemental information).

### Single-molecule optical tweezers nucleosome stability assay

To quantify the nucleosome stability at the single-molecule level, purified nucleosome with208 bp DNA containing 601 sequence was ligated with 0.8 kb and 0.5 kb lambda DNA, which were obtained by PCR using primers containing biotin or digoxigenin (BGI), respectively (Figure S10A and Table S4). These molecules allow the formation of DNA tether containing mono-nucleosome between the streptavidin (SA) bead and anti-digoxigenin (AD) bead held by the optical trap and micropipette, respectively (Figure 5A).

To measure the nucleosome outer unfolding force, inner unfolding, and refolding force, the nucleosomal DNA was incubated with 40 mM NaCl, 20 mM Tris-HCl (pH 7.5) buffer, and was pulled at 200 nm/s. To measure the inner unfolding and refolding force only, the nucleosomal DNA was incubated with 300 mM NaCl, 20 mM Tris-HCl (pH 7.5) buffer and pulled at 100 nm/s. These data were collected at 200 Hz and decimated to 40 Hz to extract the forces and the extensions.

For the extraction of the nucleosome outer wrapping and unwrapping rates, the beads were held at constant positions in the low salt buffer (5 mM NaCl, 10 mM Tris-HCl (pH 7.5)) at 2-5 pN force range. The data were collected at 1 kHz and were decimated to ~250 Hz. Transitions were determined by running a *t*-test analysis between two adjacent windows of the wrapping/unwrapping traces.

### ‘One-pot’ Assay

‘One-pot’ assay was performed the same as previously published ^31^. Briefly, ^32^P 5’-end labeled 601.2 DNA fragments with point mutations contain *Hae*III restriction digestion site at different dyad sites (Figure 2A and Table S4) were mixed in equal molar ratio for nucleosome loading, with the combination of the same cold DNA mixture (hot DNA : cold DNA= 1 : 9, molar ratio). 100 nM nucleosome was digested by *Hae*III (final concentration 1.5 U/μL) in digestion buffer (10 mM Tris, pH 7.9, 10 mM MgCl_2_, 150 mM NaCl, 20 μg/mL BSA)) at 37 °C. The reaction was stopped by adding proteinase K in the reaction stop buffer (120 mM EDTA, 0.4% SDS) and kept at 37 °C for 20 min. The final product was analyzed with 8% 1X TBE (29:1) native-PAGE and the signal was detected by Sapphire Biomolecular Imager (Azure Biosystem) and quantified with Image J.

### Structure Cryo-EM sample preparation

Nucleosome samples were loaded with 147 bp 601 DNA template and cross-linked in the 0 M reconstitution buffer with 0.15% Glutaraldehyde (Sigma) at 4 °C for 2.0 hrs. Nucleosome samples were purified with Mini Prep Cell system (Biorad). The qualified nucleosome fractions were combined and further checked by negative staining using uranyl acetate in a 3.05 mm diameter carbon film copper specimen grid. Qualified nucleosomes were then loaded in a freshly glow discharged holey carbon golden grid (Quantifoil™ R 2/2 on 300 gold mesh) (1.0-2.5 µg of nucleosomes were diluted in 40 mM NaCl reconstitution buffer in 3 μL). Movies were captured with Krios G3i cryo-TEM microscope with K3 camera (Thermo Scientific) in HKUST Biological Cryo-EM Center. Detailed microscope settings are listed in Table 1.

**Table 1.**
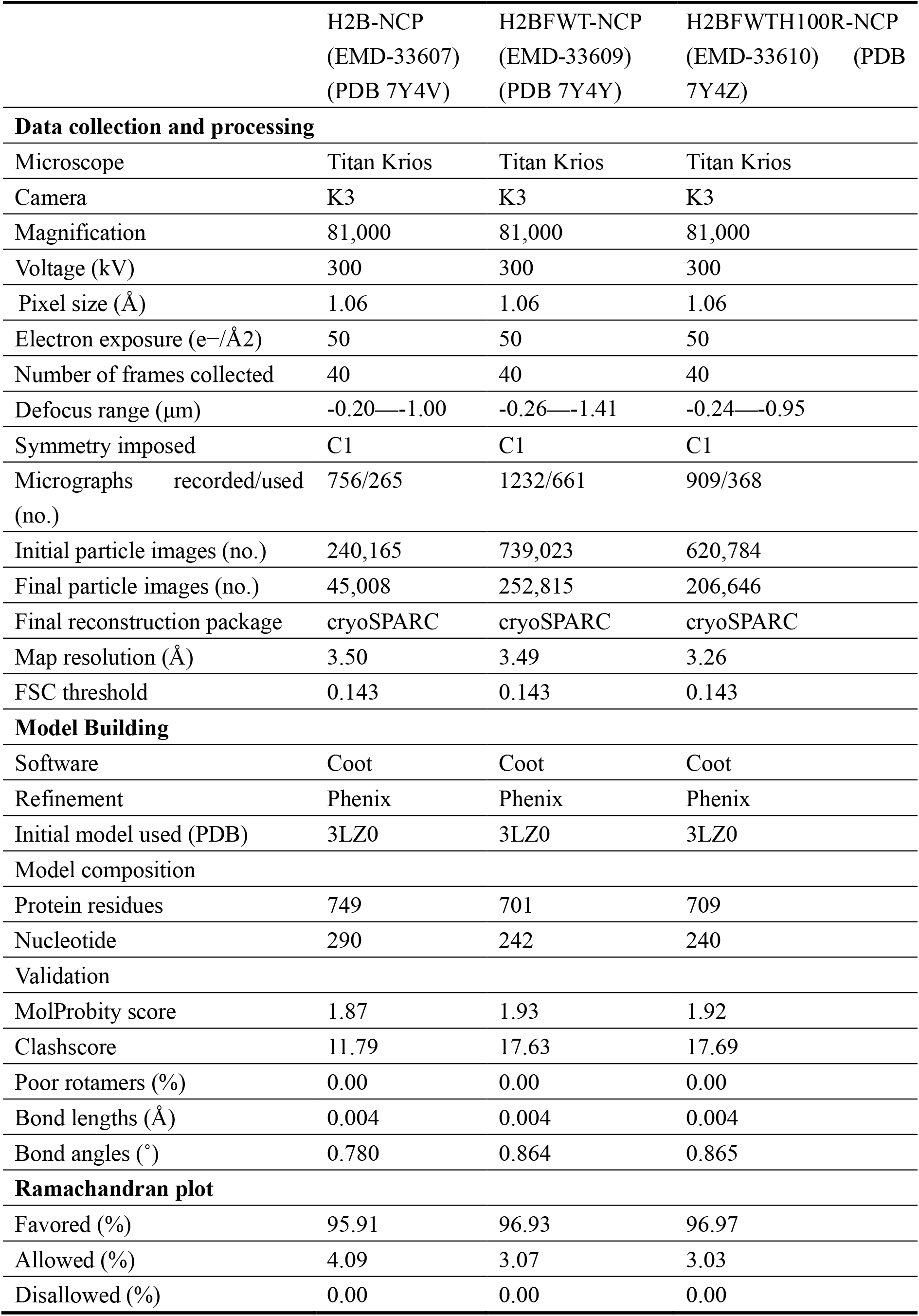
Cryo-EM data collection, refinement, and validation statistics.

### Image processing, Model building and refinement

Movies are imported into cryoSPARC (version: V2.15.0). After patch motion correction (multi)), patch CTF estimation (multi)) was performed with default settings. Manually curate exposures function was used to remove bad images. 100 micrographs were used for blob-picker to create template for template-picker. Extraction box size was set as 200 pix (1.06 Å/pix) for extract-from-micrographs. After 2D selection, ab-initio reconstruction was performed for 3D classification. Homogeneous refinement (new!) was used for mapping and the Local Resolution Estimation function was used to get the local resolution.

Template model (PDB: 3LZ0) was fitted with the density maps with Chimera (Version 1.16) and further rebuilt with Wincoot under the instruction of the user manual. Models were further refined and validated with Phenix (version 1.20-4459) real-space refinement function and comprehensive validation (cryo-EM) function. H-bond identification was performed with ChimeraX (Version 1.3) and surface electrostatics was achieved with Chimera (Version 1.16).

## Results

### H2BFWT expresses in early spermatogenesis stages, specifically in spermatogonia and primary spermatocytes in humans

As mentioned in the introduction, several groups reported that infertility patients classified as azoospermia have more mutations (or SNPs) in H2BFWT ^26–28^ (Figure S1A). Therefore, we first examined the expression pattern of the testis-specific histone variant H2BFWT in spermatogenesis by immunohistochemistry (IHC) staining using human testis (Figure 1A). Histone H4 was presented in almost all cells as expected. TH2B was mainly expressed in the later stages of spermatogenesis, including secondary spermatocyte, spermatid, and spermatozoa, which agrees with a previous study ^22^. In contrast, based on the size and position of positively-stained cells, the expression of H2BFWT was mainly limited to spermatogonia cells and primary spermatocyte cells (Figure 1A). There was little-to-no H2BFWT expression in the later stages of spermatogenesis. To confirm our observed staining patterns, we extracted the single-cell expression profiles of H2BFWT, TH2B and maker genes in human testis by reanalyzing a published single-cell transcriptome dataset (Figure S1B). The H2BFWT expression pattern matches well with STRA8 and partially with SPO11, which are markers of differentiated spermatogonia and leptotene spermatocytes, respectively ^32^, also agrees with our IHC staining result shown in Figure 1A. Furthermore, we performed an immunofluorescence staining of H2BFWT, and the H2BFWT signals can be clearly observed in the early stages of spermatogenesis (Figure 1B). Note that the H2BFWT IF signals adopt a dot-like pattern in the nucleus, suggesting that H2BFWT may localize at specific locations on the genome, which agrees with previous publication reporting that H2BFWT has co-localization with telomeres ^25^.

**Figure 1.**
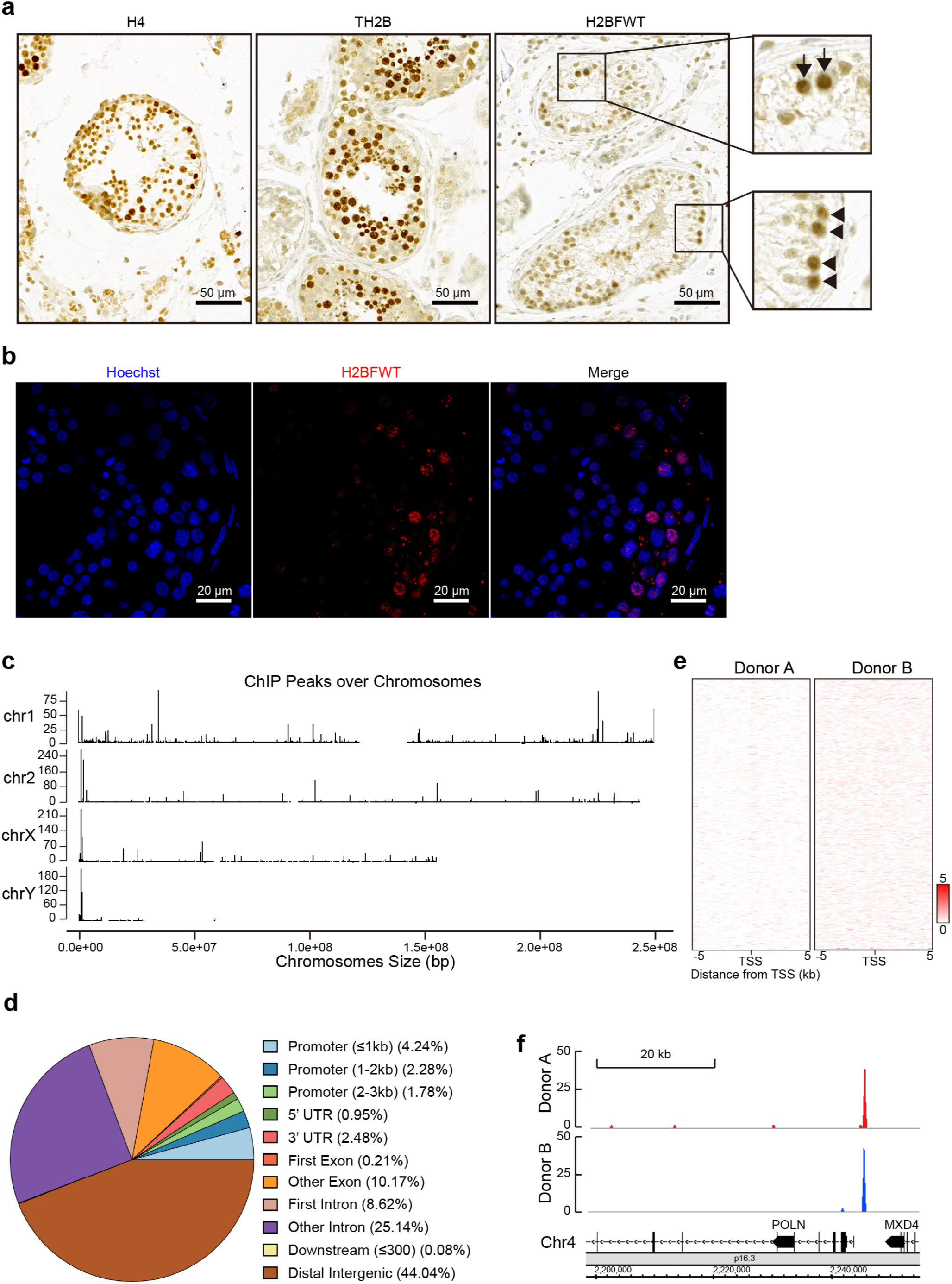
H2BFWT is specifically expressed in spermatogonia and localized in telomere/sub-telomeric regions and nearby some spermatogenesis differentiation genes in spermatocytes of human testis. (A) Immunohistochemistry staining of human testis sections with anti-H4, anti-TH2B, or anti-H2BFWT antibodies (200X). In the enlarged picture, the spermatogonia and primary spermatocyte stages are indicated with *triangles* and *arrows*, respectively. (B) Immunofluorescence staining of human testis sections with Hoechst DNA stain (blue) and anti-H2BFWT antibody (red). (C) Snapshot of H2BFWT genome wide distribution in chromosome 1, 2, X and Y. (D) H2BFWT peak genomic annotations by pie chart. (E) H2BFWT ChIP-seq profiles near TSS from two different donors. (F) The H2BFWT is enriched around the *POLN* and *HAUS3* promoter region in chromosome 4.

### H2BFWT preferentially localizes at sub-telomeric regions as well as the promoters of genes that are highly expressed in testis

To understand the detailed chromosome localization of H2BFWT in testis, we performed a ChIP-sequencing assay using human FFPE testis sections. The specificity of our homemade H2BFWT antibody was verified using ChIP-qPCR with stable Flag-H2BFWT HeLa cells. We found that our anti-H2BFWT antibody could detect a qPCR pattern highly similar to the anti-Flag antibody control (Figure S2).

After analyzing the H2BFWT-ChIP libraries (GEO: GSE206260) from human FFPE testis samples, we found that H2BFWT was highly enriched at the ends of most chromosomes (Figure 1C and S3). This telomere or sub-telomere enrichment is consistent with previous reports that GFP-H2BFWT foci overlapped with telomere structures found in cultured cell imaging ^25^. When the ChIP peaks were annotated, H2BFWT is slightly enriched in the genic region, although we found no particular H2BFWT enrichment pattern near TSS or TES regions (Figure 1D, E, S4A and S4B). H2BFWT localization on the testicular genome seems to be quite sporadic except in telomeric or sub-telomeric regions (Figure S3 and S4C). Although the genome-wide localization of H2BFWT may not be related to the global gene transcription in testis, we found that H2BFWT preferentially localized around some specific testicular high expression genes (Figure S5A), and the promoter/enhancer regions of *HDAC2, TMX4, POLN*, and *HAUS3*, in which the later two which shares a 5’ exon (Figure 1F) ^33^. Interestingly, we noted that *POLN* and *HAUS3* express at the same stages as H2BFWT in the testicular single-cell expression analysis (Figure S1B and S5B). We then checked whether H2BFWT affects the expression of *POLN* and *HAUS3* using a Dox-inducible Flag-H2BFWT HeLa cell line. We found that H2BFWT expression enhances *POLN* and *HAUS3* gene expression in a H2BFWT-dose-dependent manner (Figure S5C). Our results suggest that H2BFWT is mostly localized at the telomere and sub-telomeric regions and it is also involved in driving the expression of a few testicular high expression genes.

### H2BFWT destabilizes the nucleosome and its H100R SNP further reduces nucleosomal stability

Since H2BFWT seems to act as a gene-specific expression activator in testis, we checked whether this activating effect is due to the formation of a destabilized nucleosome by H2BFWT. Furthermore, as mentioned in the introduction, the H2BFWT H100R SNP is more frequently observed in infertile patients, therefore, we are also curious how this particular substitution changes the nucleosome structure and/or stability. First, nucleosomes reconstituted *in vitro* were incubated with different salt buffers to quantify their stability (Figure S6). With increasing salt concentrations, free DNA and non-nucleosomal DNA-histone complexes gradually appear, and their relative amounts keep increasing in both H2B and H2BFWT nucleosome samples (Figure S6A). However, the amounts and relative percentage of free DNA exhibit larger increases for the H2BFWT nucleosome than those for the H2B nucleosome (Figure S6B). This result indicates that the H2BFWT nucleosome is less stable than the H2B nucleosome. However, no statistical difference was observed between the wild-type H2BFWT and the H2BFWTH100R nucleosomes (Figure S6C and D). We hypothesized that the salt stability assay might not be sensitive enough to distinguish the effect of a single amino acid substitution. Therefore, we further performed a ‘one-pot’ nucleosome digestion assay (Figure 2). We found that the H2BFWT nucleosome is more sensitive to digestion by the restriction enzyme *Hae*III along the entire nucleosomal regions (d0 - d7) (Figure 2B, 2C and Figure S7). Particularly, the H2BFWT nucleosome was digested more at the d_3_ and d_5_ regions where the H2BFWT N-terminus interacts with DNA. Furthermore, we found that the H2BFWTH100R nucleosome is even more sensitive to digestion by *Hae*III when compared to the H2BFWT nucleosome, even though it is only a point mutation, it appears to exert a destabilizing effect to the whole nucleosome over all nucleosomal regions (Figure 2B, 2D and Figure S7).

**Figure 2.**
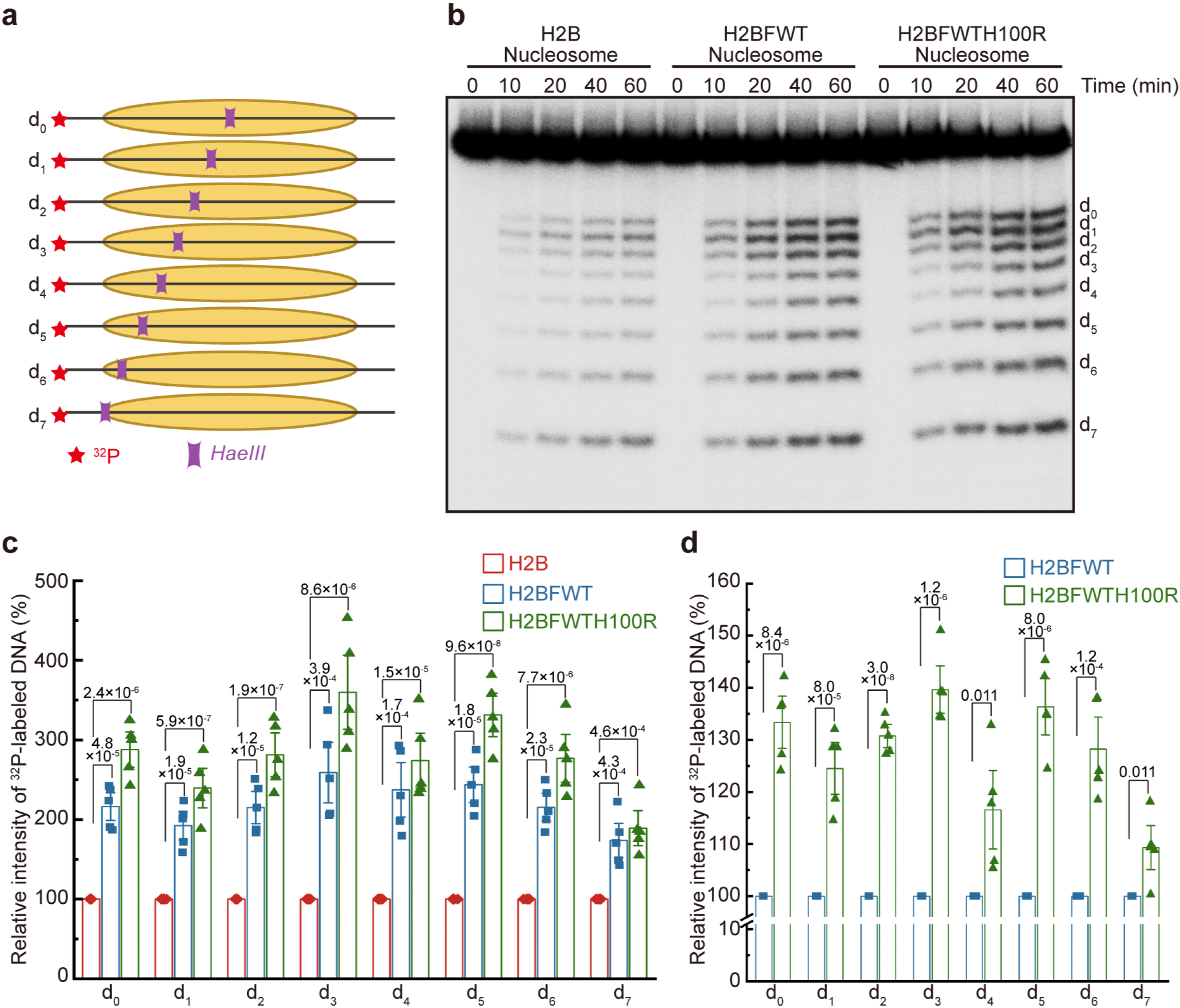
H2BFWT and H2BFWT H100R nucleosomes are more sensitive to restriction enzyme digestion in One-pot Assay. (A) Nucleosomal DNA accessibility was examined with ‘one-pot’ assay. Single-end ^32^P-labeled DNA containing *Hae*III restriction site at the indicated positions was used in equal molar ratio during nucleosomes reconstitution. (B) A representative image of *Hae*III digestion in ‘one-pot’ assay. The ^32^P-labeled DNA fragments after the digestion were detected by phosphor imaging. (C) The histogram of relative band intensities of digested ^32^P-labeled DNA fragments in H2BFWT and H2BFWTH100R nucleosomes compared with H2B nucleosomes. Data are represented as mean ± SEM from independent experiments (n = 5). (D) The histogram of relative band intensities of digested ^32^P-labeled DNA fragments in H2BFWTH100R nucleosomes compared with H2BFWT nucleosomes. Data are represented as mean ± SEM from independent experiments (n = 5).

### H2BFWT weakens the interactions between H2B-H4 interfaces in nucleosomes

To further understand the destabilization effects of H2BFWT and its H100R mutation, we determined the cyro-EM structure of the human H2BFWT nucleosome (Figure 3 and Table 1, PDB: 7Y4Y). In total ~739,000 particles were picked from micrographs of H2BFWT nucleosomes and classified into several distinct classes (Figure S8). Further analyses selected ~253,000 particles of homogeneous H2BFWT nucleosome, leading to a final EM map with 3.49 Å resolution of the H2BFWT nucleosome (Figure 3, S8, and Table 1). Our structure analysis shows that, while the overall architecture of H2BFWT nucleosome is similar to the canonical nucleosome (Figure 3 and Table 1, PDB: 7Y4V), the H2BFWT nucleosome undergoes some notable conformational and structural rearrangements. Compared to the canonical nucleosome, we found that the DNA ends of the H2BFWT nucleosome become floppy, indicating a more dynamic structure in the relevant regions (Figure 3A-C). Moreover, the H3 N-terminus, the H2A N-terminus and the H2BFWT N-terminus all have less extension in the H2BFWT nucleosome compared with the canonical control (Figure 3D). In contrast, the C-terminus of H2BFWT in the H2BFWT nucleosome has a longer extension than that of H2B in the H2B nucleosome (Figure 3D). The H3 N-terminus, H2A N-terminus, and H2B/H2BFWT N-terminus are all positively charged whereas H2BFWT C-terminus is negatively charged according to the surface electrostatics map (Figure 3D). These differences between H2BFWT nucleosome and H2B nucleosome make the H2BFWT nucleosome structure more relaxed and less stable.

**Figure 3.**
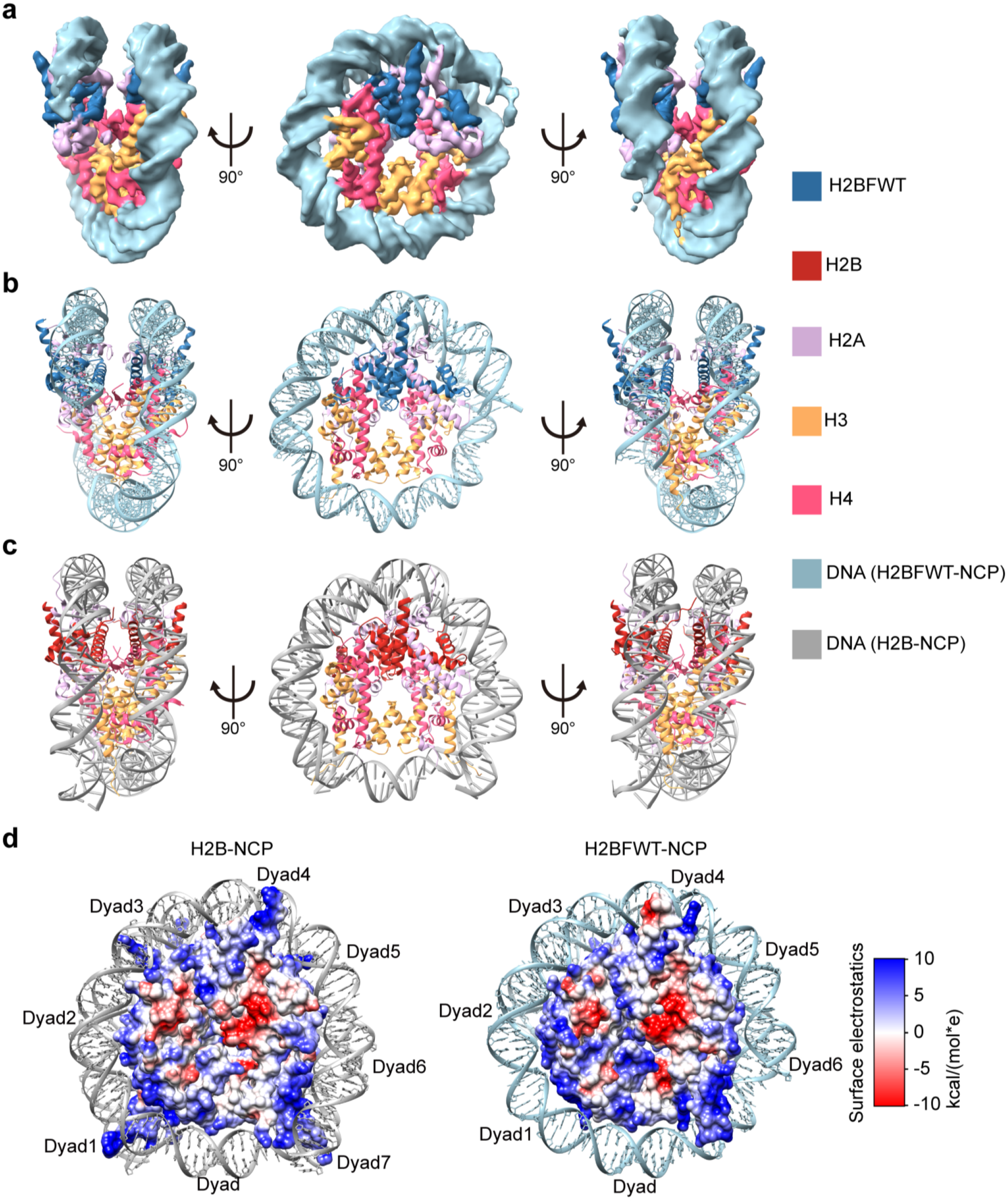
Cryo-EM structures of H2BFWT and canonical nucleosomes. (A) Cryo-EM density map of the H2BFWT nucleosome at 3.49 Å resolution. Disc view (middle) and gyre views (left and right). (B) Atomic model of the H2BFWT nucleosome in disc view (middle), and gyre views (left and right). (C) Atomic model of the H2B nucleosome in disc view (middle), and gyre views (left and right). (D) Surface electrostatic (red, negative; blue, positive; potential display levels were between −10 and 10 kcal/(mol*e)) on H2B octamer and H2BFWT octamer in the presence of DNA.

The structure of H2BFWT in nucleosomes is overall similar to that of H2B, with the exception that the length of the αC helix was increased, and the positions of loop 1 and loop 3 were different (Figure 4A). In loop 3, the H2B E105 which constitutes part of the acidic patch between H2A and H2B, is replaced by a neutral amino acid glutamine (Q) at the residue 126 in H2BFWT (Figure 4B). The acidic patch is an important region for chromatin formation and protein-nucleosome interactions such as 53BP1 and HMGN2 ^1,34,35^. H2B Loop 1 is mainly involved in the intra-nucleosomal interactions with DNA. In the canonical nucleosome, Y40, Y42, and S56 form hydrogen bonds with DNA. However, equivalent residues in H2BFWT, Y61 and R63 cannot form hydrogen bonds, and only R77 can form a single H-bond with the DNA backbone (Figure 4C and Table S1). Another interesting part is the L1 loop in H2A. The H2A L1 loop is important for the H2A/H2B dimer-dimer interaction. In the H2BFWT nucleosome, however, the H2A L1 loop position is altered, and a new H-bond is formed between two H2A L1 loops (Figure 4D). Furthermore, H2BFWT significantly reduces the number of interactions between the H2A-H2BFWT dimer and the H3-H4 tetramer, as sites available for H-bond formation are decreased between H2BFWT and H4 (Table S2). Apart from these, the H2BFWT nucleosome also has less H3-H4 tetramer-DNA interactions than the H2B nucleosome (Figure 4E, Table S3).

**Figure 4.**
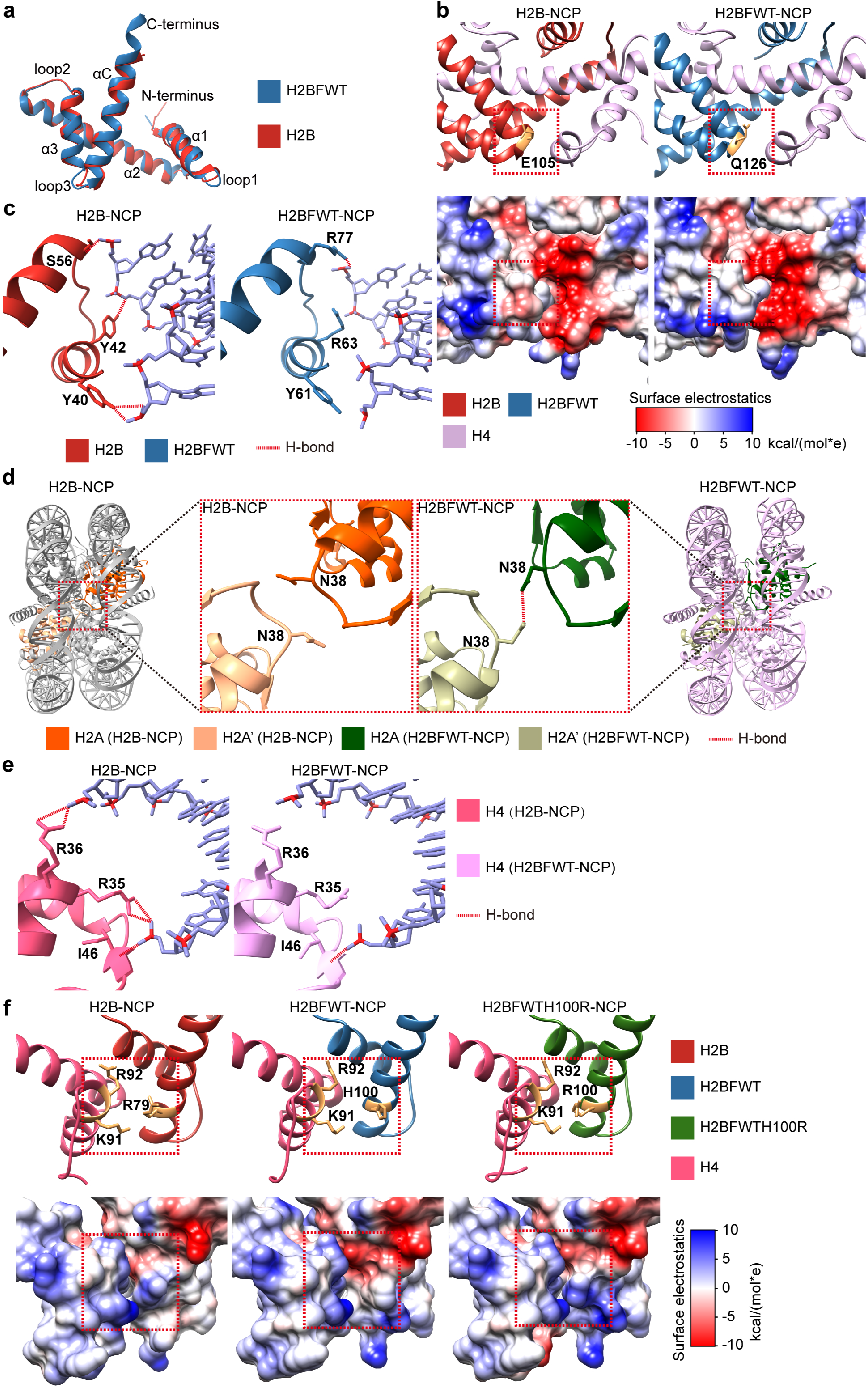
H2BFWT destabilizes nucleosomal structure through weakening of interactions between H2BFWT and DNA, H2BFWT and H4, as well as H3-H4 tetramer and DNA. (A) Conformational comparison between H2B and H2BFWT models within the nucleosome. (B) Close-up view of the acidic patch region of H2B and H2BFWT atomic models (upper) and surface electrostatics (lower). Red frame indicates the acidic patch region with differences in the H2B and the H2BFWT nucleosomes. (C) Close-up view of the loop 1 region and nearby DNA in the H2B and the H2BFWT nucleosomes. (D) Close-up view of the H2A L1 loop regions in the H2B and the H2BFWT nucleosomes. (E) Close-up view of H4 α1 region and nearby DNA in the H2B and the H2BFWT nucleosomes. (F) Close-up view of the interaction regions between H2B/H2BFWT/H2BFWTH100R and H4 displaying as atomic models (upper) and surface electrostatics (lower).

We also successfully resolved the structure of the H2BFWTH100R nucleosome at 3.26 Å resolution (Figure S9 and Table 1, PDB: 7Y4Z). H2BFWTH100 is located at the interface region between H2BFWT and H4. Our structure analysis indicates that the conformations of this region are very similar in the canonical H2B and the H2BFWT nucleosomes, however, the H100R substitution modifies the surface electrostatics to positive which crashes with the positively charged H4K91 in addition to changing the side chain angle of R100 (Figure 4F). This result well-explain why the stability of the H2BFWTH100R nucleosome is further reduced.

### H2BFWTH100R increases the nucleosome unwrapping rate which further reduces the nucleosome stability from the H2BFWT nucleosome

To quantify and dissect the nucleosome destabilization effect of H2BFWT, we measured the H2BFWT nucleosome stability using several optical tweezers single-molecule assays (Figure 5). The nucleosomes to be tested were loaded onto and ligated with Digoxin and Biotin tagged DNA fragments for optical trapping (Figure 5A). As shown in Figure 5B and consistent with previous publications, there are two nucleosome unwrapping events as the force/distance increases, namely the outer and inner rips. The outer rip represents the interaction between H2A/H2B dimer and DNA, and the inner rip represents the interaction between H3/H4 tetramer and DNA ^36,37^.

**Figure 5.**
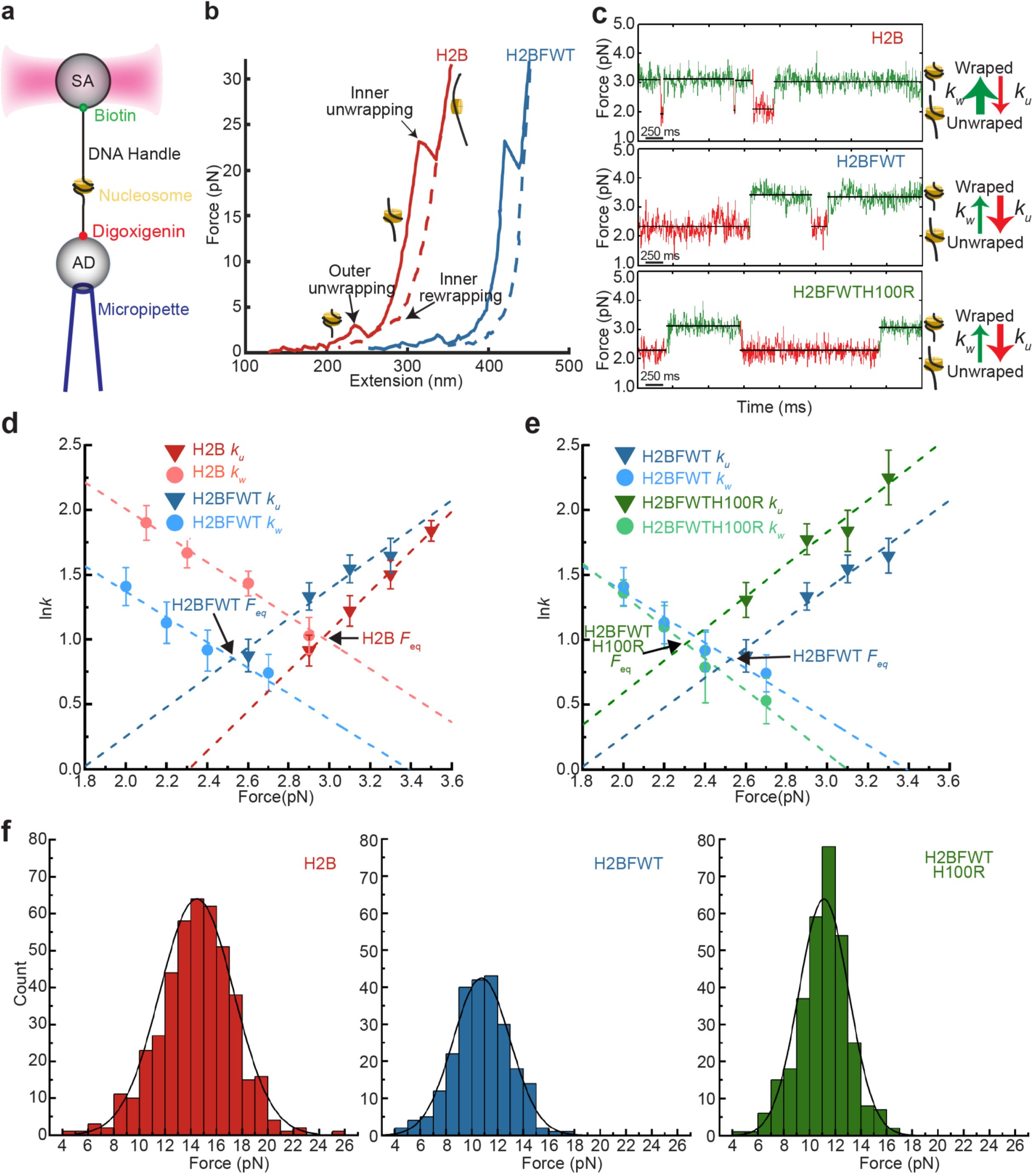
H2BFWT nucleosomes possess increased unwrapping rate and decreased rewrapping rate which signifies destabilizations of the outer region of the nucleosome. (A) Geometry of the single-molecule optical tweezers nucleosome assay. (B) Example traces of nucleosome pulling of H2BFWT and the H2B nucleosomes. (C) Examples of unwrapping and rewrapping in the outer region of the H2B (upper panel), the H2BFWT (middle panel), and the H2BFWTH100R (bottom panel) nucleosomes. (D) The unwrapping rate (*k*_*u*_) and rewrapping rate (*k*_*w*_) of the H2BFWT and the H2B nucleosomes plotted in natural logarithm against applied force. (E) The unwrapping rate (*k*_*u*_) and rewrapping rate (*k*_*w*_) of the H2BFWT and the H2BFWTH100R nucleosomes plotted in natural logarithm against applied force. (F) The inner unwrapping force distribution of the H2B, the H2BFWT, and the H2BFWTH100R nucleosomes.

To examine the effects of H2BFWT and H2BFWTH100R on the H2A/H2BFWT dimer and nucleosomal DNA interaction (i.e. the outer rip) in details, a nucleosome optical tweezers hopping experiment was performed under low salt concentration (5 mM) by holding a nucleosome tether at a fixed distance (Figure 5C). During this experiment, when the position of the nucleosome was kept at a suitable location or the force was held within an appropriate range, the nucleosome will dynamically transition between the outer wrapped state (green) and the outer unwrapped state (red) (Figure 5C-E). These dynamic changes are nucleosome outer hoppings. The hopping force range is determined to be 2.0–4.0 pN for the H2B nucleosome, and 1.5-3.5 pN for the H2BFWT and the H2BFWTH100R nucleosomes (Figure 5D, 5E, and Table 2). To analyze outer hopping in more detail, the wrapping rate (*k*_*w*_) and the unwrapping rate (*k*_*u*_) were calculated at different holding forces by measuring the time of the unwrapped and wrapped states. The *k*_*w*_ and *k*_*u*_ rates are defined as the reciprocal of the time of a given state (i.e. the unwrapped state or the wrapped state). When *k*_*w*_ and *k*_*u*_ rates were plotted against force in natural logarithm, it was found that with increasing force, the *k*_*u*_ rates will increase and *k*_*w*_ rates will decrease for all three nucleosomes (i.e. H2B, the H2BFWT and the H2BFWTH100R) (Figure 5D and 5E).

**Table 2.** Summary of the optical tweezers data of the H2B, the H2BFWT and the H2BFWT100R nucleosomes.

| $k_u$ | | | | | | $k_w$ | | | | | |
| --- | --- | --- | --- | --- | --- | --- | --- | --- | --- | --- | --- |
| H2B |  | H2BFWT |  | H2BFWT100R |  | H2B |  | H2BFWT |  | H2BFWT100R |  |
| Force<br>(pN) | $k_u$<br>(n)* | Force<br>(pN) | $k_u$<br>(n)* | Force<br>(pN) | $k_u$<br>(n)* | Force<br>(pN) | $k_w$<br>(n)* | Force<br>(pN) | $k_w$<br>(n)* | Force<br>(pN) | $k_w$<br>(n)* |
| 2.90<br>(2.70-3.00) | 2.5±0.3<br>(106) | 2.60<br>(2.53-2.70) | 2.4±0.3<br>(43) | 2.60<br>(2.58-2.73) | 3.7±0.5<br>(60) | 2.10<br>(1.95-2.20) | 6.7±0.9<br>(97) | 2.00<br>(1.85-2.08) | 4.1±0.6<br>(60) | 2.00<br>(1.90-2.10) | 3.9±0.5<br>(82) |
| 3.10<br>(3.00-3.20) | 3.4±0.4<br>(134) | 2.90<br>(2.73-3.00) | 3.8±0.4<br>(120) | 2.90<br>(2.73-3.01) | 5.9±0.8<br>(125) | 2.30<br>(2.20-2.50) | 5.3±0.6<br>(184) | 2.20<br>(2.08-2.25) | 3.1±0.5<br>(81) | 2.20<br>(2.10-2.30) | 3.0±0.4<br>(49) |
| 3.30<br>(3.20-3.40) | 4.5±0.5<br>(113) | 3.10<br>(3.00-3.15) | 4.7±0.5<br>(62) | 3.10<br>(3.01-3.15) | 6.3±1.0<br>(44) | 2.60<br>(2.50-2.75) | 4.2±0.4<br>(154) | 2.40<br>(2.25-2.55) | 2.5±0.4<br>(113) | 2.40<br>(2.30-2.50) | 2.2±0.6<br>(45) |
| 3.50<br>(3.40-3.60) | 6.3±0.5<br>(139) | 3.30<br>(3.15-3.63) | 5.2±0.7<br>(129) | 3.30<br>(3.16-3.60) | 9.5±2.0<br>(54) | 2.90<br>(2.75-3.00) | 2.8±0.4<br>(154) | 2.70<br>(2.55-3.20) | 2.1±0.3<br>(76) | 2.70<br>(2.50-3.00) | 1.7±0.3<br>(31) |

(n)\* means number of events happened at each force range.

The equilibrium force (*F*_*eq*_), which is the applied force where *k*_*w*_ and *k*_*u*_ are equal, is 2.97 pN for the H2B nucleosome, and is decreased to 2.54 pN for the H2BFWT nucleosome. More interestingly, *F*_*eq*_ further decreased to 2.27 pN in the H2BFWTH100R nucleosome (Figure 5D, 5E and Table 3). The free energy cost *ΔG*^*0*^ of the outer rip in the H2B nucleosome was calculated as 20.5 kJ/mol. For the H2BFWT nucleosome, the *ΔG*^*0*^ decreased by around 40% to 12.84 kJ/mol, indicating that the H2A/H2BFWT dimer has much weaker interactions with nucleosomal DNA. Moreover, the least stable H2BFWT100R nucleosome possesses the lowest *ΔG*^*0*^ at 9.36 kJ/mol (Table 3).

**Table 3.** *F*_*eq*_ and energy barriers (*ΔG*^*0*^) of the H2B, the H2BFWT and the H2BFWTH100R nucleosomes.

| | $F_{eq}$ (pN) | Energy Barrier<br>$\Delta G^0$ (kJ/mol) |
| --- | --- | --- |
| H2B | 2.97 | 20.46 |
| H2BFWT | 2.54 | 12.84 |
| H2BFWTH100R | 2.27 | 9.36 |

Furthermore, we also found that the inner unwrapping forces were lower in the H2BFWT and the H2BFWTH100R nucleosomes in high salt condition (Figure 5F and Figure S10C), however, no change was observed on the inner rewrapping forces (Figure S10B and C). This result agrees with the fact that the number of H-bond between the H3-H4 tetramer and DNA was reduced in the H2BFWT nucleosome (Table S3). To dispel the possibility that hexasomes originated from spontaneous nucleosome dissociation at diluted conditions such as those suitable for optical tweezers experiments would skew the wrapping/unwrapping force measurements, we checked the integrities of the nucleosomes with native PAGE electrophoresis. We discovered that only two bands (free DNA and the nucleosome bands) existed with no signs of hexasomes or other undesirable histone-DNA complexes even when the nucleosomes were incubated at the same dilution level as the optical tweezers experiment for a prolonged period (Figure S10D).

### H2BFWT nucleosome poses a weaker barrier for Pol II transcription elongation than the canonical nucleosome

To understand how the weaker H2BFWT nucleosome enhances gene expression, we performed a Pol II transcription elongation assay with H2B- or H2BFWT-containing nucleosomes (Figure 6A, 6B). As previously reported, the nucleosomal passage efficiencies of Pol II (i.e. the Run-off percentage) increase along with KCl concentrations in the reaction mixtures (Figure 6C) ^38^. By quantifying the Run-off percentage, we found that Pol II has an easier time transcribing through the H2BFWT nucleosome than the H2B nucleosome, with the effect being more pronounced at higher salt conditions (Figure 6C and 6D). This trend stays the same for both yeast and mammalian RNA polymerases (Figure 6 and Figure S11). The nucleosome entry region (+15nt) represents the interaction between H2A/H2B dimer and DNA (Figure 6C and 6E). Around this region, H2BFWT and H2BFWTH100R nucleosomes appear to pose much fewer barriers for Pol II transcription which manifests as fewer and fainter gel bands (Figure 6C and 6E). Altogether, results of the transcription assays suggest that H2A/H2BFWT dimer has weaker interaction with the nucleosomal DNA, which is consistent with our optical tweezers assays and cryo-EM structural models. Coupled with the observed H2BFWT genomic localization, our study suggests that H2BFWT likely functions to enhance spermatogenesis-related gene transcription in spermatogonia by decreasing Pol II transcription barriers at specific testicular high expression genes.

**Figure 6.**
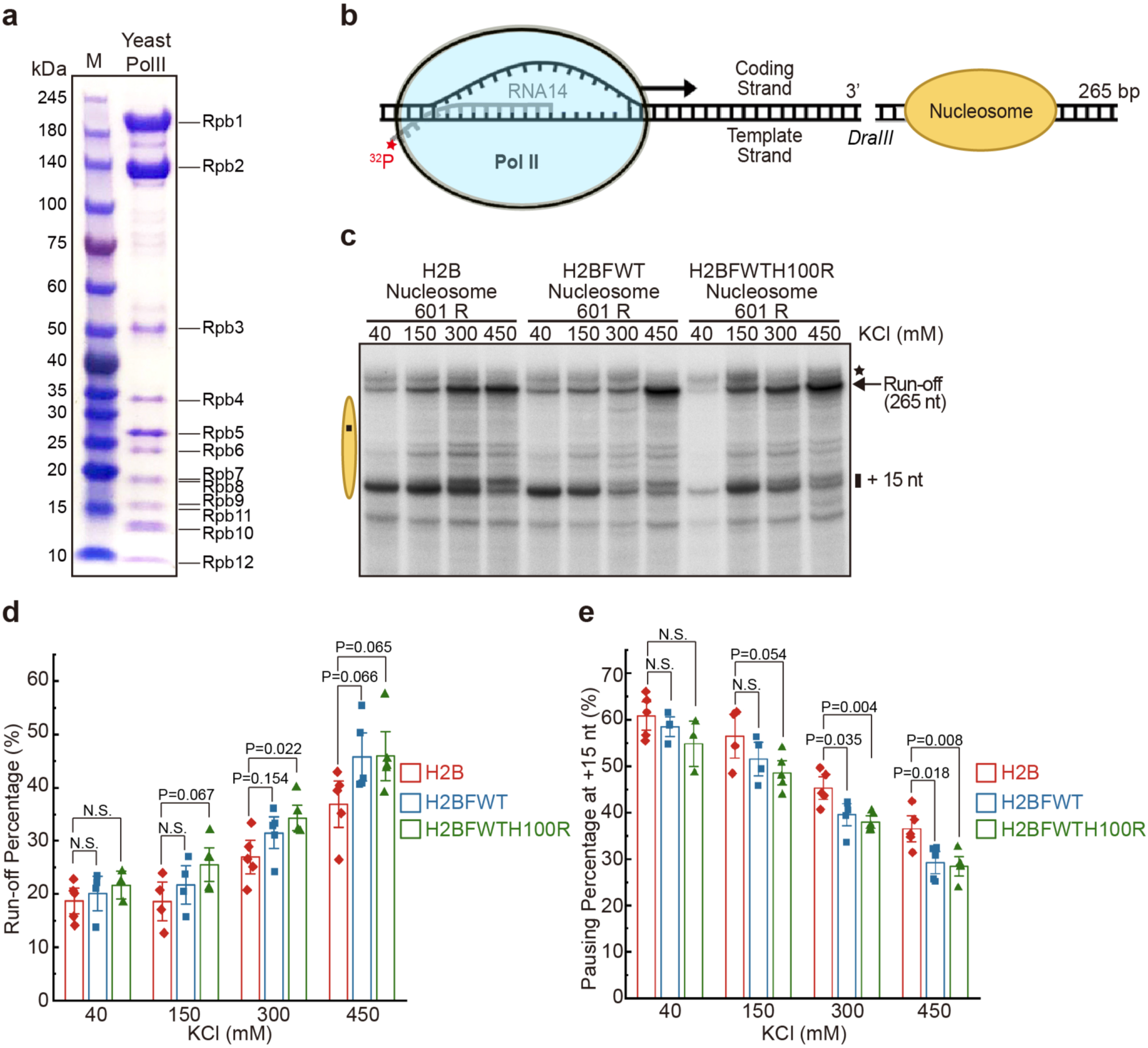
Pol II transcribes more efficiently through H2BFWT nucleosomes. (A) The representative SDS-PAGE gel image of the purified yeast Pol II. (B) Schematic diagram of the Pol II *in vitro* transcription model on nucleosomal DNA templates. H2B, H2BFWT or H2BFWTH100R nucleosomes were ligated with the TEC as the transcription templates in the experiments. (C) Nucleosome templates were transcribed in the presence of the indicated concentrations of KCl. Arrow indicates the position of Run-off transcripts. The position of the nucleosome on the template is indicated by the oval (nucleosome dyad region is indicated with a black square). +15 nt region is labeled on the right side. Star indicates the secondary structure of Run-off products. (D) The percentage of run-off transcripts was quantified under different salt conditions. Error bars represent SEM in repeating experiments (n=5). (E) The relative percentage of pausing at +15 nt position. Error bars represent SEM in repeating experiments (n=5).

## Discussion

Primate-specific histone H2B variant H2BFWT is localized in differentiating spermatogonia and early spermatocytes and is enriched in the telomeric region. The H2BFWT localization is correlated with the activation of testicular gene expression by means of nucleosome destabilization. Because of the weaker interactions between H2A-H2BFWT dimer and DNA or H3-H4 tetramer, Pol II experiences significantly less transcriptional resistance. As a result, expressions of H2BFWT-enriched genes are enhanced. More interestingly, our results show that even though canonical histones remain the same in terms of primary sequences, the shape of H2A L1 loop and H3-H4 tetramer-DNA interaction are altered in the H2BFWT nucleosome, demonstrating that the structural effect of H2BFWT extends across the nucleosome. The H2A L1 loop is an important region mediating the H2A/H2B dimer-dimer interactions. A change in L1 loop conformation is also observed in the transcriptionally repressive macroH2A nucleosome, which renders the L1-L1 interface to make it less flexible and more hydrophobic, resulting in a more stable nucleosome ^39^. In another example, even though the overall structure of the H2A.Z nucleosome is similar to the canonical nucleosome, interactions between the L1 loops of H2A.Zs are strengthened and thus the H2A.Z histone octamer is likely more stable ^40^. Swapping the H2B to H2BFWT may have a similar effect as histone H2A variant replacements. Moreover, H2BFWT lacks the H2BK120 equivalent residue where it is known to be ubiquitinated at actively transcribing genes which prevent the dissociation of H2A/H2B dimers during Pol II transcription ^41^. This is another possible mechanism that H2BFWT may enhance transcription activities.

Another interesting residuelocation is the H2BFWTQ126 residue, which is the spatial equivalent of H2BE105. They are a part of the nucleosome acidic patch, which consists of six H2A (E56, E61, E64, D90, E91 and E92) and two H2B (E105 and E113) residues. The acidic patch is well known to be involved in chromatin formation and protein-histone interactions, which include several chromatin remodeling factors and histone PTM modifying enzymes ^42^. It is reported that acidic patch mutations can alter the function of chromatin remodelers ^43^. The amino acid substitution from the negatively charged glutamate (E) in H2BE105 to the neutral glutamine (Q) in H2BFWTQ126 likely weakens the acidic patch. This may, in turns, alter the structure and interacting proteins of H2BFWT-containing chromatin, and possibly even the histone PTM patterns. It would not be a surprise if the presence of H2BFWT creates a unique and necessary genomic environment specific to the early spermatogenesis stage.

From previous papers and our own ChIP-sequencing analysis of human testis, we found that H2BFWT preferentially localizes in the telomeric/subtelomeric region. It is possible that this localization may assist the regulation of appropriate telomere dynamics during spermatogenesis. It is known that the telomere goes through dynamical changes during spermatogenesis ^44^ and that the length of telomere in sperm is linked to human fertility ^45^. It is a real possibility that H2BFWT may also be involved in telomere boundary control during histone-protamine transition in spermatogenesis.

H2BFWT is only expressed in testis and is specific to primates. Compared to other tissues, the testis has the highest number of expressed genes ^46,47^. Furthermore, the testis is a highly transcriptionally active tissue, and it is also the only tissue that is capable of producing new genes in male ^48,49^. In fact, a wave of DNA substitution mutagenesis was observed in early spermatogonia after germline stem cells enter spermatogenesis. Through this process, germline stem cells keep dividing for self-population renewal and to produce fate-determined spermatogonia cells for male gamete production. Although the DNA repair machinery is upregulated during meiosis, DNA-repairing activities drop-off sharply at later stages of spermatogenesis, it is proposed that this is a mechanism by which some mutations remain unfixed and subsequently transferred to the next generation which accounts for the appearance of *de novo* genes ^50^. This may well explain why H2BFWT is only present in primates and its expression is limited to testis.

As mentioned in the introduction section, there are two SNPs of H2BFWT related to infertility. The SNP at the 5’UTR regions (−9C>T) introduces a new start codon ATG at the −10 position and causes a frameshift. Multiple studies asserted that no H2BFWT protein can be produced ^26–28^. It is also reported that this SNP is strongly-associated with azoospermia ^27^. Therefore, evidence supports the idea that H2BFWT has important functions in spermatogenesis. The SNP at 368A>G, which substitutes the amino acid at position 100 from histidine (H) to arginine (R) (H2BFWTH100R), is associated with a less-severe phenotype of oligospermia, in which sperm is produced but its number is low ^27^. In this study, we found that the arginine substitution changes the interaction between H2BFWTH100R and H4, resulting in exacerbated nucleosome destabilization. This further destabilized H2BFWTH100R nucleosome very likely disrupts the regulation of H2BFWT-controlled testicular genes to some extent, however, since H2BFWTH100R can still form nucoeomses, the H2BFWT-related gene regulatory network may only be partially perturbed. This well explains the reported association of H2BFWTH100R (368A>G) with the milder phenotype of oligospermia instead of azoospermia like the −9C>T nonsense SNP.

Finally, from our structural model, H2BFWTH100 is facing the H4K91 and H4R92 residues. Histone H4K91 can be post-translationally modified in several different ways. It is highly acetylated in the preleptotene stage ^51^, which roughly coincides with the expression timing of H2BFWT. In addition to acetylation, H4K91 ubiquitination was detected in testis ^52^. H4R92 is also reported to be mono-methylated in mouse spermatogenic cells and human sperms ^53^. It is, therefore, possible that H2BFWTH100R changes the surface electrostatics and the spacing between H2BFWTH100 and H4K91, which indirectly affects the post-translational modifications of H4K91 and H4R92. But because the functional roles of H4K91 and H4R92 PTMs in spermatogenesis remain currently unclear, it would be interesting to investigate how this H100R substitution in H2BFWT changes gene expression patterns and if it exerts any effect on telomere length maintenance during spermatogenesis in humans.

## Acknowledgement

This work was supported by the Research Grants Council of the Hong Kong SAR [C7009-20G] to YL and TI, the Hong Kong Innovation & Technology Commission Innovation and Technology Fund [MHP/033/20] and National Science Foundation of China [32170548] to TI. We thank Prof. Jie Qiao from the Peking University to provide the single cell sequencing TPM data and Prof. Patrick Cramer from Max Planck Institute for Biophysical Chemistry to provide the hexahistidine-tagged Pol II *S. cerevisiae* strain. The EM dataset was collected at the Biological Cryo-EM Center, generously supported by a donation from the Lo Kwee Seong Foundation, at HKUST.

## References

1. Luger, K., Mäder, A.W., Richmond, R.K., Sargent, D.F. & Richmond, T.J. Crystal structure of the nucleosome core particle at 2.8 Å resolution. Nature 389, 251–260 (1997).

2. Luger, K., Dechassa, M.L. & Tremethick, D.J. New insights into nucleosome and chromatin structure: an ordered state or a disordered affair? Nature reviews Molecular cell biology 13, 436–447 (2012).

3. Yuan, G. & Zhu, B. Histone variants and epigenetic inheritance. Biochimica et Biophysica Acta (BBA)-Gene Regulatory Mechanisms 1819, 222–229 (2012).

4. Long, M. et al. A novel histone H4 variant H4G regulates rDNA transcription in breast cancer. Nucleic Acids Research 47, 8399–8409 (2019).

5. Ausio, J. Histone variants—the structure behind the function. Briefings in Functional Genomics 5, 228–243 (2006).

6. Martire, S. & Banaszynski, L.A. The roles of histone variants in fine-tuning chromatin organization and function. Nature reviews Molecular cell biology 21, 522–541 (2020).

7. Rathke, C., Baarends, W.M., Awe, S. & Renkawitz-Pohl, R. Chromatin dynamics during spermiogenesis. Biochimica et Biophysica Acta (BBA)-Gene Regulatory Mechanisms 1839, 155–168 (2014).

8. Ausio, J., Zhang, Y. & Ishibashi, T. Histone variants and posttranslational modifications in spermatogenesis and infertility. in Epigenetic Biomarkers and Diagnostics 479–496 (Elsevier, 2016).

9. Tachiwana, H. et al. Structural basis of instability of the nucleosome containing a testis-specific histone variant, human H3T. Proceedings of the National Academy of Sciences 107, 10454–10459 (2010).

10. Tachiwana, H., Osakabe, A., Kimura, H. & Kurumizaka, H. Nucleosome formation with the testis-specific histone H3 variant, H3t, by human nucleosome assembly proteins in vitro. Nucleic acids research 36, 2208–2218 (2008).

11. Ueda, J. et al. Testis-specific histone variant H3t gene is essential for entry into spermatogenesis. Cell reports 18, 593–600 (2017).

12. Urahama, T. et al. Histone H3. 5 forms an unstable nucleosome and accumulates around transcription start sites in human testis. Epigenetics & chromatin 9, 1– 16 (2016).

13. Schenk, R., Jenke, A., Zilbauer, M., Wirth, S. & Postberg, J. H3. 5 is a novel hominid-specific histone H3 variant that is specifically expressed in the seminiferous tubules of human testes. Chromosoma 120, 275–285 (2011).

14. Eirín‐López, J.M., Ishibashi, T. & Ausió, J. H2A. Bbd: a quickly evolving hypervariable mammalian histone that destabilizes nucleosomes in an acetylation‐independent way. The FASEB Journal 22, 316–326 (2008).

15. Chadwick, B.P. & Willard, H.F. A novel chromatin protein, distantly related to histone H2A, is largely excluded from the inactive X chromosome. The Journal of cell biology 152, 375–384 (2001).

16. Bao, Y. et al. Nucleosomes containing the histone variant H2A. Bbd organize only 118 base pairs of DNA. The EMBO journal 23, 3314–3324 (2004).

17. Soboleva, T.A. et al. A unique H2A histone variant occupies the transcriptional start site of active genes. Nature structural & molecular biology 19, 25–30 (2012).

18. Nekrasov, M., Soboleva, T.A., Jack, C. & Tremethick, D.J. Histone variant selectivity at the transcription start site: H2A. Z or H2A. Lap1. Nucleus 4, 431– 437 (2013).

19. Soboleva, T.A. et al. A new link between transcriptional initiation and pre-mRNA splicing: The RNA binding histone variant H2A. B. PLoS genetics 13, e1006633 (2017).

20. Tolstorukov, M.Y. et al. Histone variant H2A. Bbd is associated with active transcription and mRNA processing in human cells. Molecular cell 47, 596–607 (2012).

21. Padavattan, S. et al. Structural and functional analyses of nucleosome complexes with mouse histone variants TH2a and TH2b, involved in reprogramming. Biochem Biophys Res Commun 464, 929–35 (2015).

22. Montellier, E. et al. Chromatin-to-nucleoprotamine transition is controlled by the histone H2B variant TH2B. Genes & development 27, 1680–1692 (2013).

23. Shinagawa, T. et al. Disruption of Th2a and Th2b genes causes defects in spermatogenesis. Development 142, 1287–1292 (2015).

24. Churikov, D. et al. Novel human testis-specific histone H2B encoded by the interrupted gene on the X chromosome. Genomics 84, 745–756 (2004).

25. Boulard, M. et al. The NH2 tail of the novel histone variant H2BFWT exhibits properties distinct from conventional H2B with respect to the assembly of mitotic chromosomes. Molecular and cellular biology 26, 1518–1526 (2006).

26. Lee, J. et al. Functional polymorphism in H2BFWT‐5′ UTR is associated with susceptibility to male infertility. Journal of cellular and molecular medicine 13, 1942–1951 (2009).

27. Ying, H.-q., Scott, M.B. & Zhou-Cun, A. Relationship of SNP of H2BFWT gene to male infertility in a Chinese population with idiopathic spermatogenesis impairment. Biomarkers 17, 402–406 (2012).

28. Rafatmanesh, A., Nikzad, H., Ebrahimi, A., Karimian, M. & Zamani, T. Association of the c.‐9C> T and c. 368A> G transitions in H2BFWT gene with male infertility in an Iranian population. Andrologia 50, e12805 (2018).

29. Fanelli, M. et al. Pathology tissue–chromatin immunoprecipitation, coupled with high-throughput sequencing, allows the epigenetic profiling of patient samples. Proceedings of the National Academy of Sciences 107, 21535–21540 (2010).

30. Cejas, P. et al. Chromatin immunoprecipitation from fixed clinical tissues reveals tumor-specific enhancer profiles. Nature medicine 22, 685–691 (2016).

31. Jing, Y. et al. Semisynthesis of site-specifically succinylated histone reveals that succinylation regulates nucleosome unwrapping rate and DNA accessibility. Nucleic acids research 48, 9538–9549 (2020).

32. Wang, M. et al. Single-cell RNA sequencing analysis reveals sequential cell fate transition during human spermatogenesis. Cell stem cell 23, 599-614. e4 (2018).

33. Takata, K.-i. et al. Conserved overlapping gene arrangement, restricted expression, and biochemical activities of DNA polymerase ν (POLN). Journal of Biological Chemistry 290, 24278–24293 (2015).

34. Wilson, M.D. et al. The structural basis of modified nucleosome recognition by 53BP1. Nature 536, 100–103 (2016).

35. Kato, H. et al. Architecture of the high mobility group nucleosomal protein 2-nucleosome complex as revealed by methyl-based NMR. Proceedings of the National Academy of Sciences 108, 12283–12288 (2011).

36. Bintu, L. et al. Nucleosomal elements that control the topography of the barrier to transcription. Cell 151, 738–749 (2012).

37. Hall, M.A. et al. High-resolution dynamic mapping of histone-DNA interactions in a nucleosome. Nature structural & molecular biology 16, 124– 129 (2009).

38. Wan, Y.C.E. et al. Cancer-associated histone mutation H2BG53D disrupts DNA–histone octamer interaction and promotes oncogenic phenotypes. Signal transduction and targeted therapy 5, 1–4 (2020).

39. Chakravarthy, S. et al. Structural characterization of the histone variant macroH2A. Molecular and cellular biology 25, 7616–7624 (2005).

40. Suto, R.K., Clarkson, M.J., Tremethick, D.J. & Luger, K. Crystal structure of a nucleosome core particle containing the variant histone H2A. Z. Nature structural biology 7, 1121–1124 (2000).

41. Wyce, A. et al. H2B ubiquitylation acts as a barrier to Ctk1 nucleosomal recruitment prior to removal by Ubp8 within a SAGA-related complex. Molecular cell 27, 275–288 (2007).

42. Markert, J. & Luger, K. Nucleosomes meet their remodeler match. Trends in Biochemical Sciences 46, 41–50 (2021).

43. Dao, H.T. & Pham, L.T. Acidic patch histone mutations and their effects on nucleosome remodeling. Biochemical Society Transactions 50, 907–919 (2022).

44. Fice, H.E. & Robaire, B. Telomere dynamics throughout spermatogenesis. Genes 10, 525 (2019).

45. Turner, S. & Hartshorne, G. Telomere lengths in human pronuclei, oocytes and spermatozoa. MHR: Basic science of reproductive medicine 19, 510–518 (2013).

46. Parisi, M. et al. A survey of ovary-, testis-, and soma-biased gene expression in Drosophila melanogasteradults. Genome biology 5, 1–18 (2004).

47. Soumillon, M. et al. Cellular source and mechanisms of high transcriptome complexity in the mammalian testis. Cell reports 3, 2179–2190 (2013).

48. Long, M., Betrán, E., Thornton, K. & Wang, W. The origin of new genes: glimpses from the young and old. Nature Reviews Genetics 4, 865–875 (2003).

49. Neme, R. & Tautz, D. Fast turnover of genome transcription across evolutionary time exposes entire non-coding DNA to de novo gene emergence. elife 5, e09977 (2016).

50. Witt, E., Benjamin, S., Svetec, N. & Zhao, L. Testis single-cell RNA-seq reveals the dynamics of de novo gene transcription and germline mutational bias in Drosophila. Elife 8, e47138 (2019).

51. Getun, I.V. et al. Functional roles of acetylated histone marks at mouse meiotic recombination hot spots. Molecular and cellular biology 37, e00942–15 (2017).

52. Lai, F. et al. Identification of histone modifications reveals a role of H2b monoubiquitination in transcriptional regulation of dmrt1 in Monopterus albus. International journal of biological sciences 17, 2009 (2021).

53. Luense, L.J. et al. Comprehensive analysis of histone post-translational modifications in mouse and human male germ cells. Epigenetics & chromatin 9, 1–15 (2016).

