## Supplemental Figures and Tables for "Testis-specific H2BFWT disrupts nucleosome integrity through reductions of DNA-histone interactions"

Figure S1 Ding *et al.*,

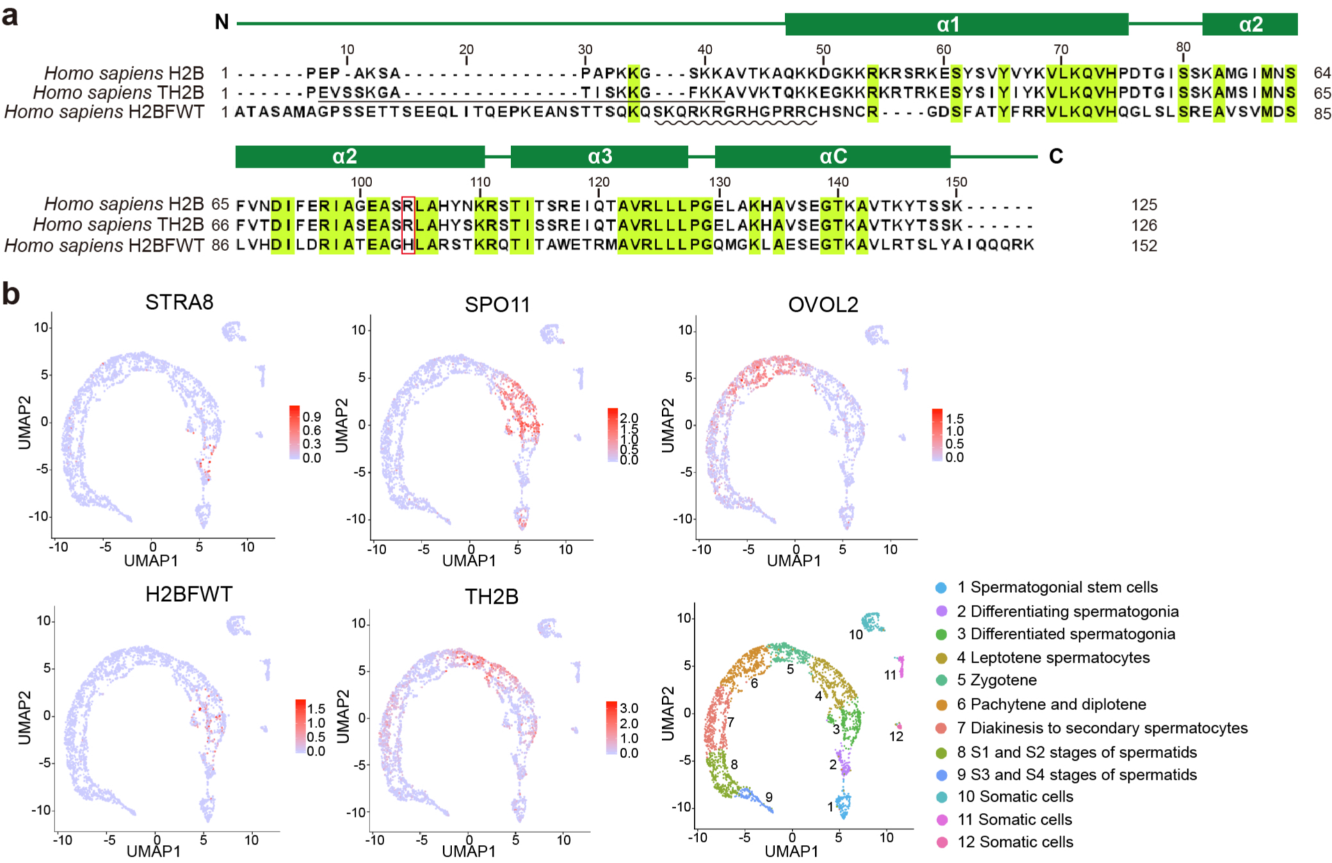

Figure S2 Ding *et al.*,

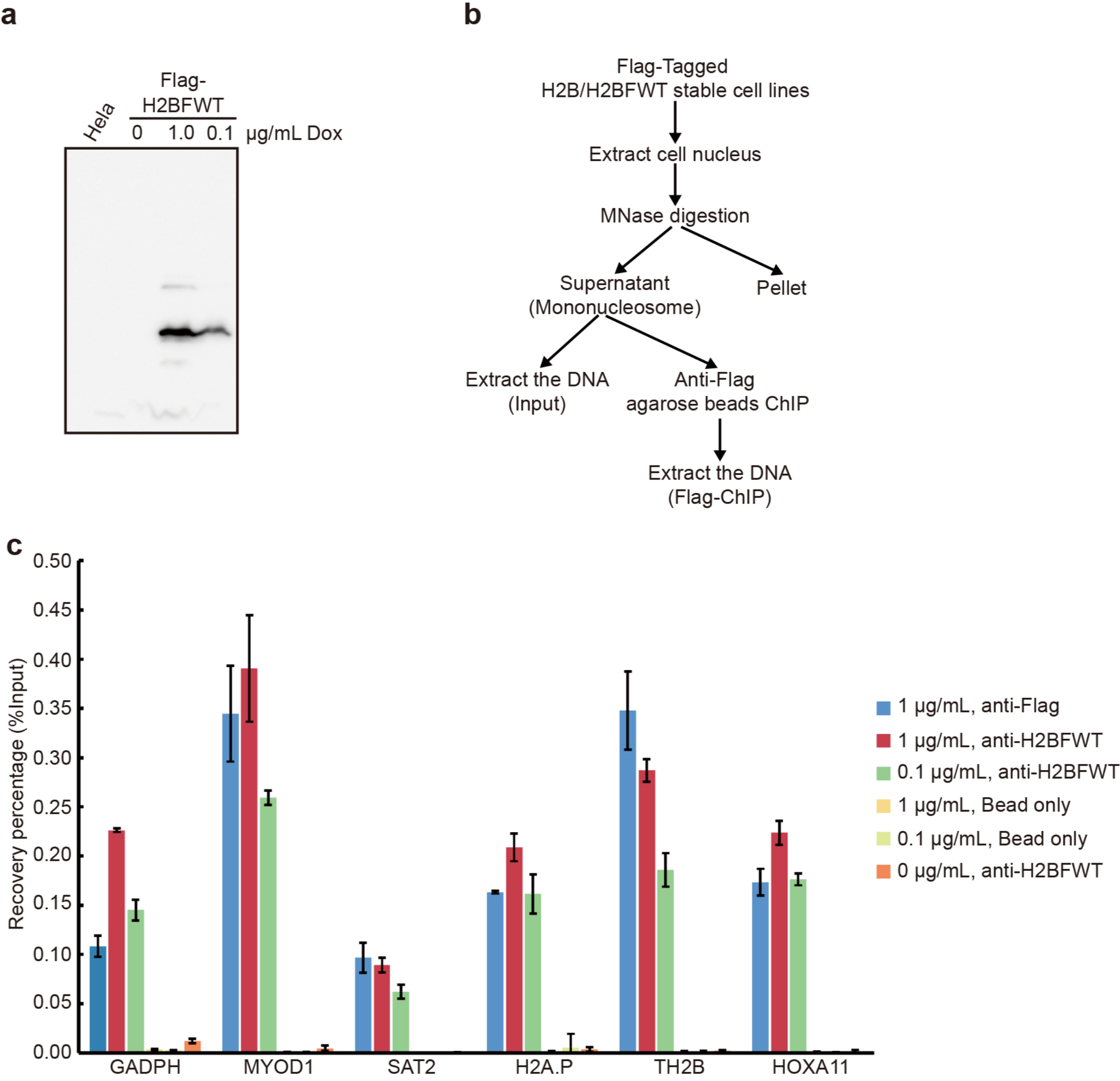

Figure S3 Ding *et al.*,

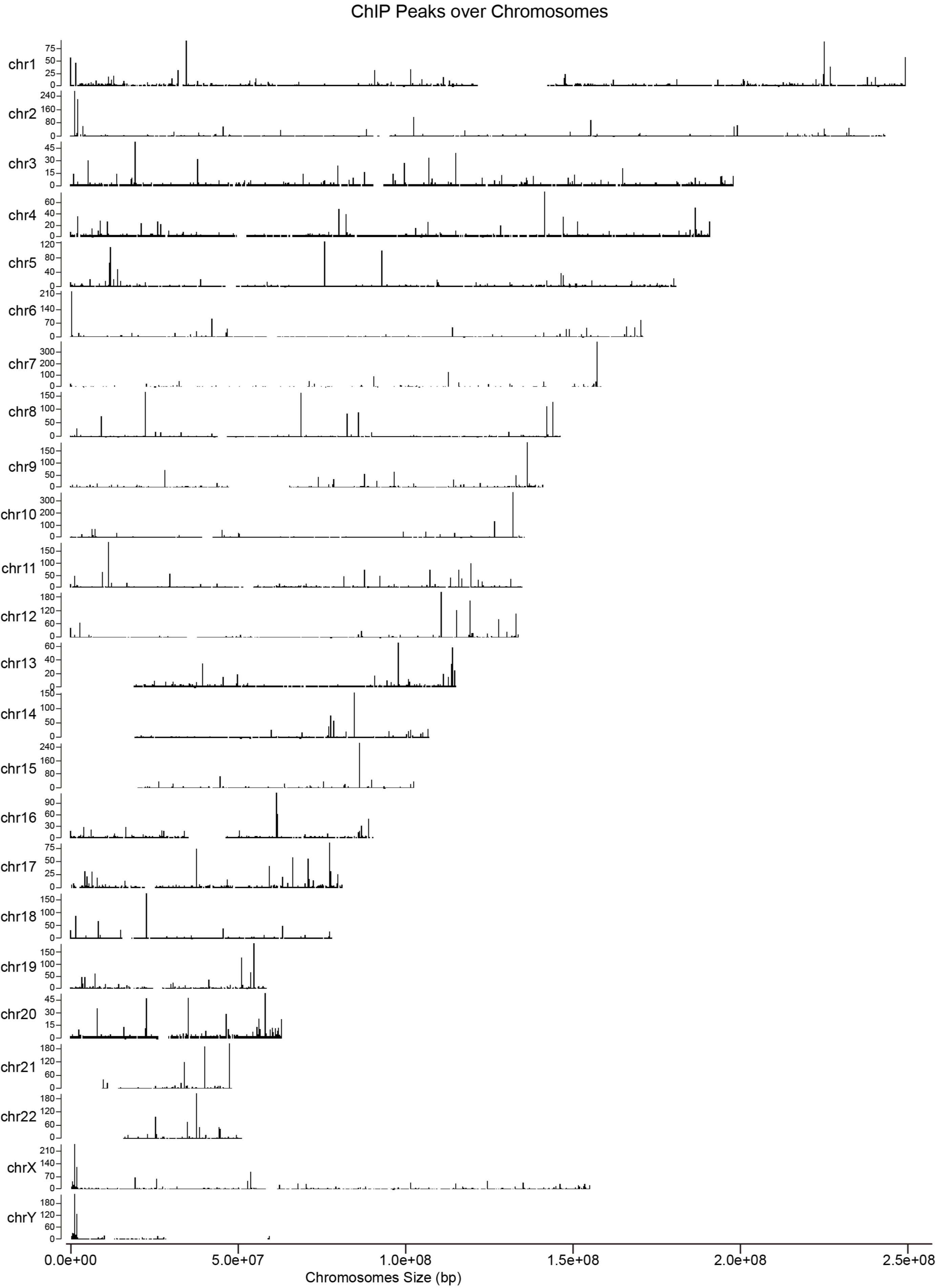

Figure S4 Ding *et al.*,

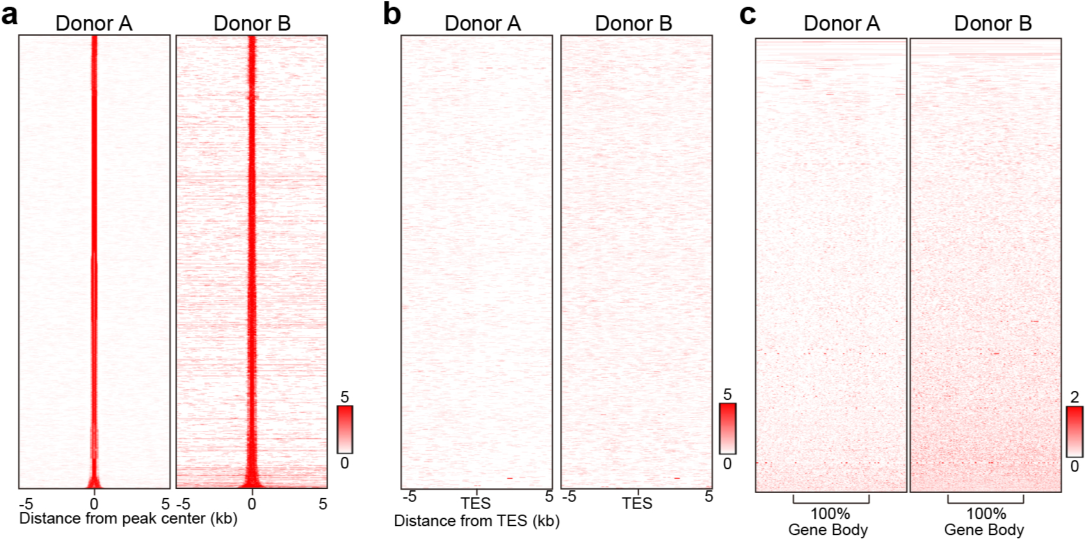

Figure S5 Ding *et al.*,

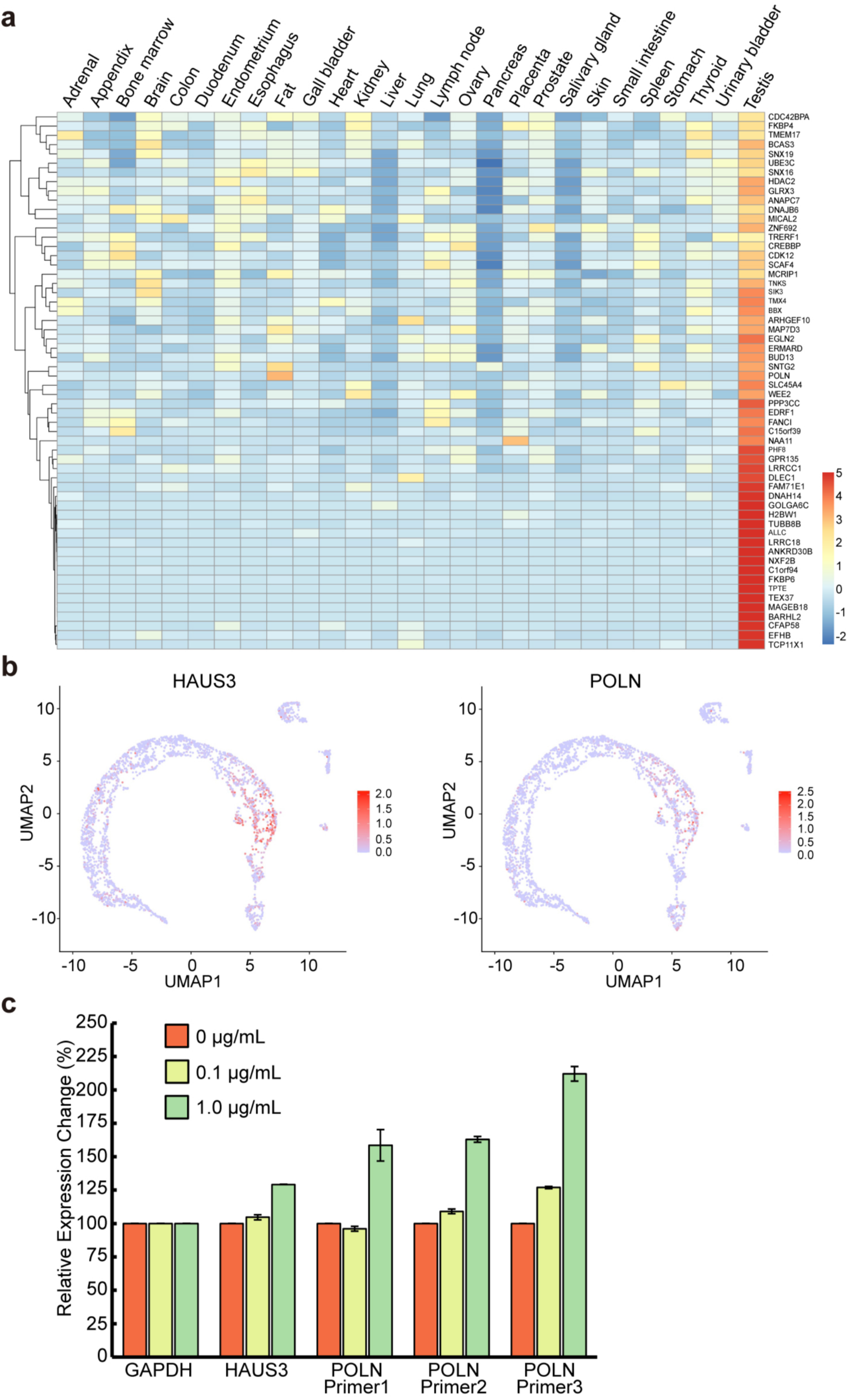

Figure S6 Ding *et al.*,

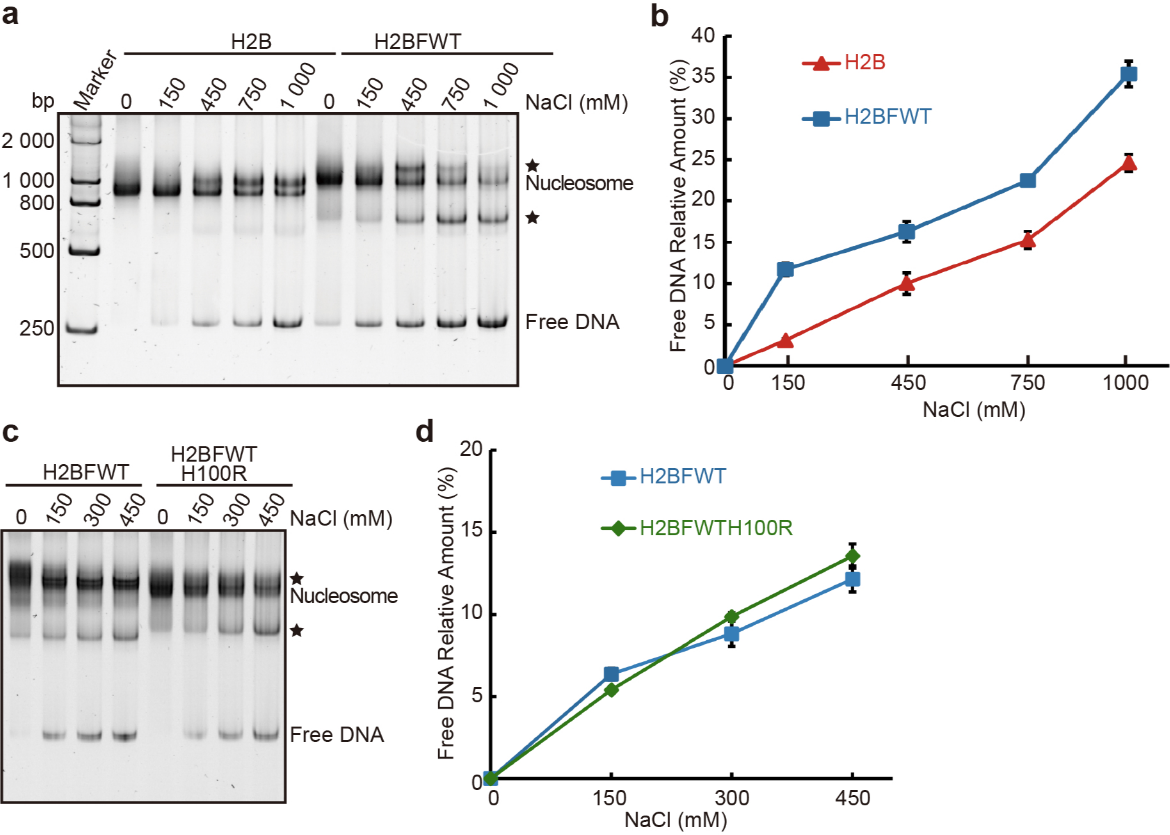

Figure S7 Ding *et al.*,

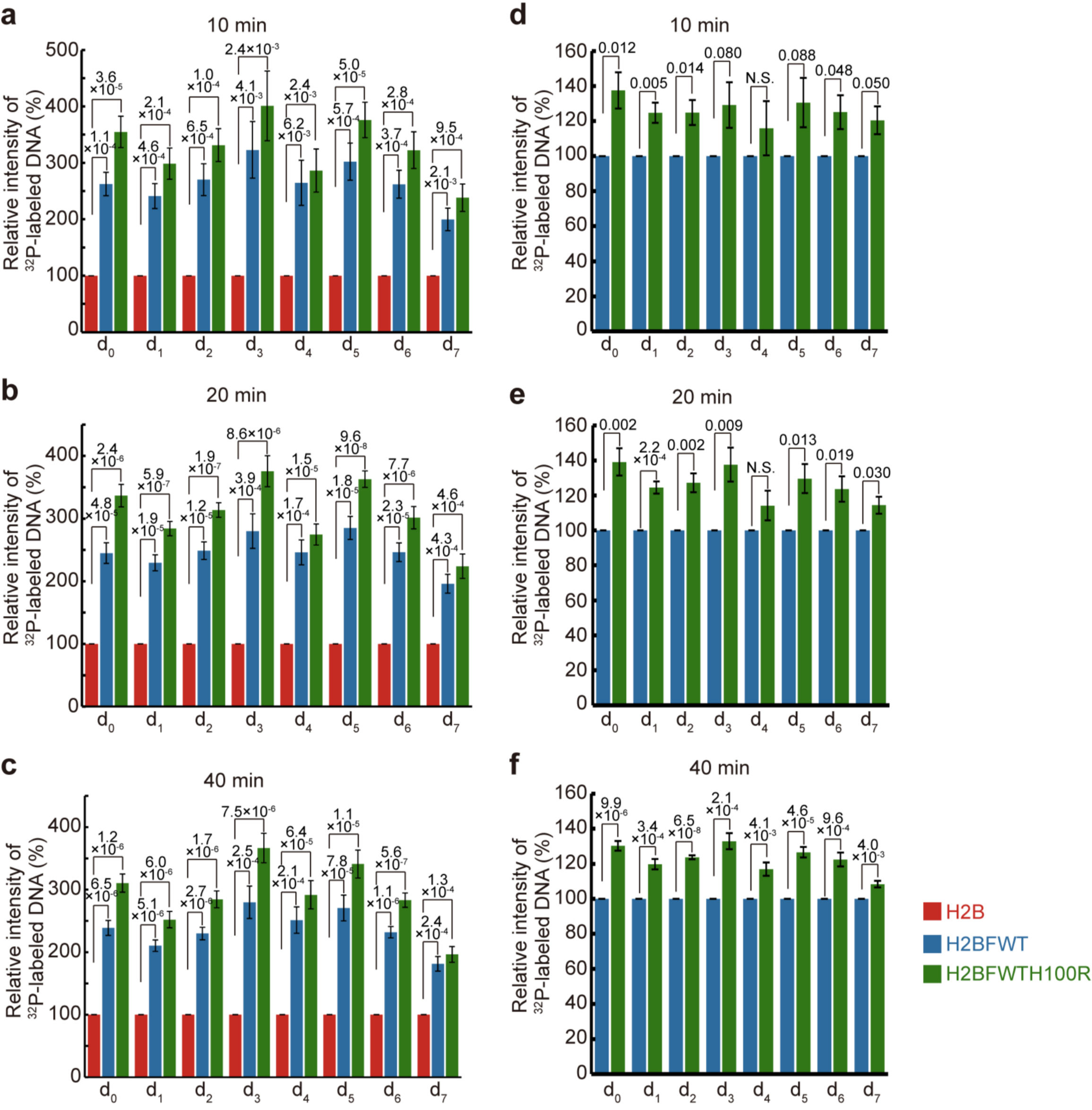

Figure S8 Ding *et al.*,

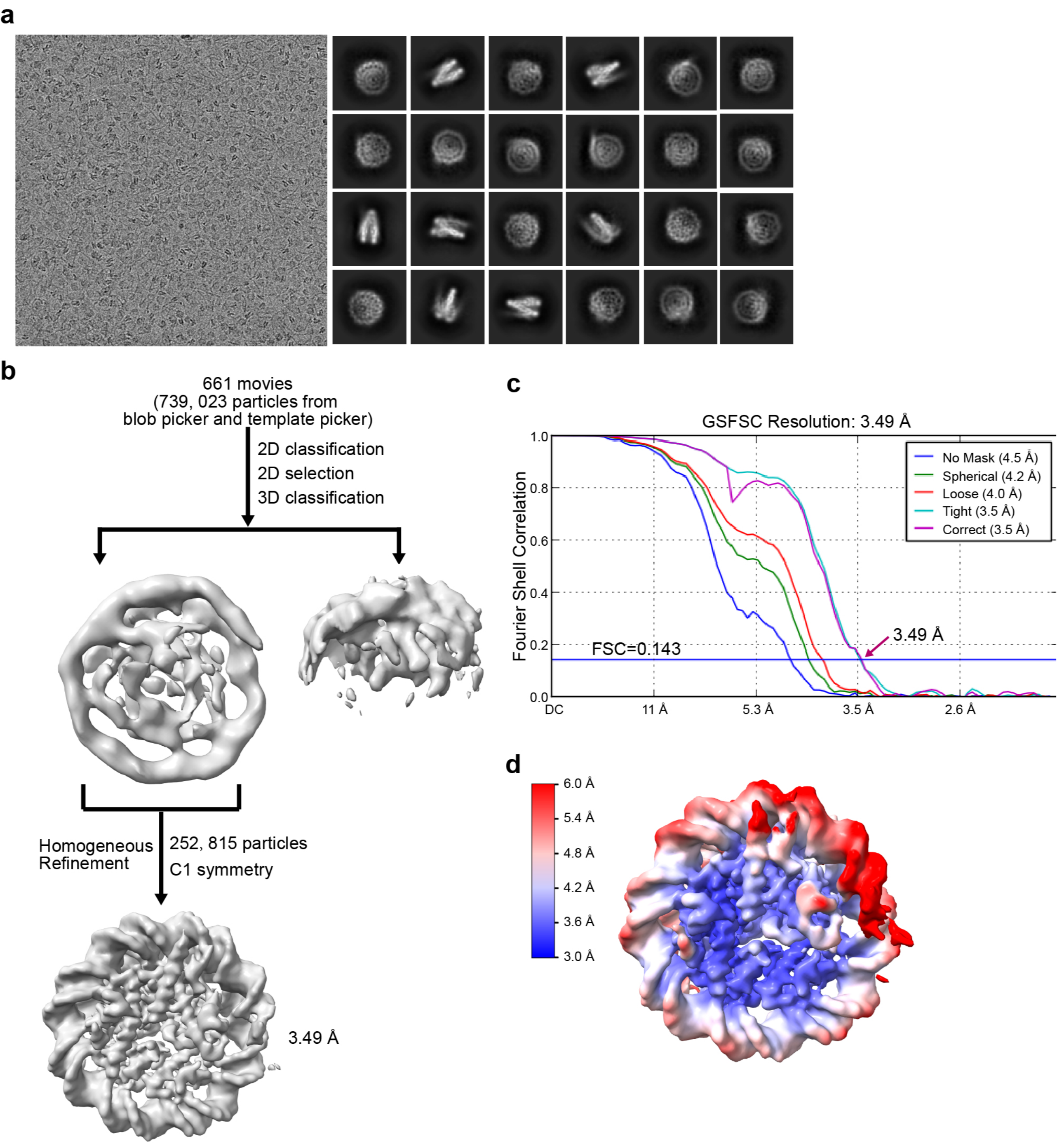

Figure S9 Ding *et al.*,

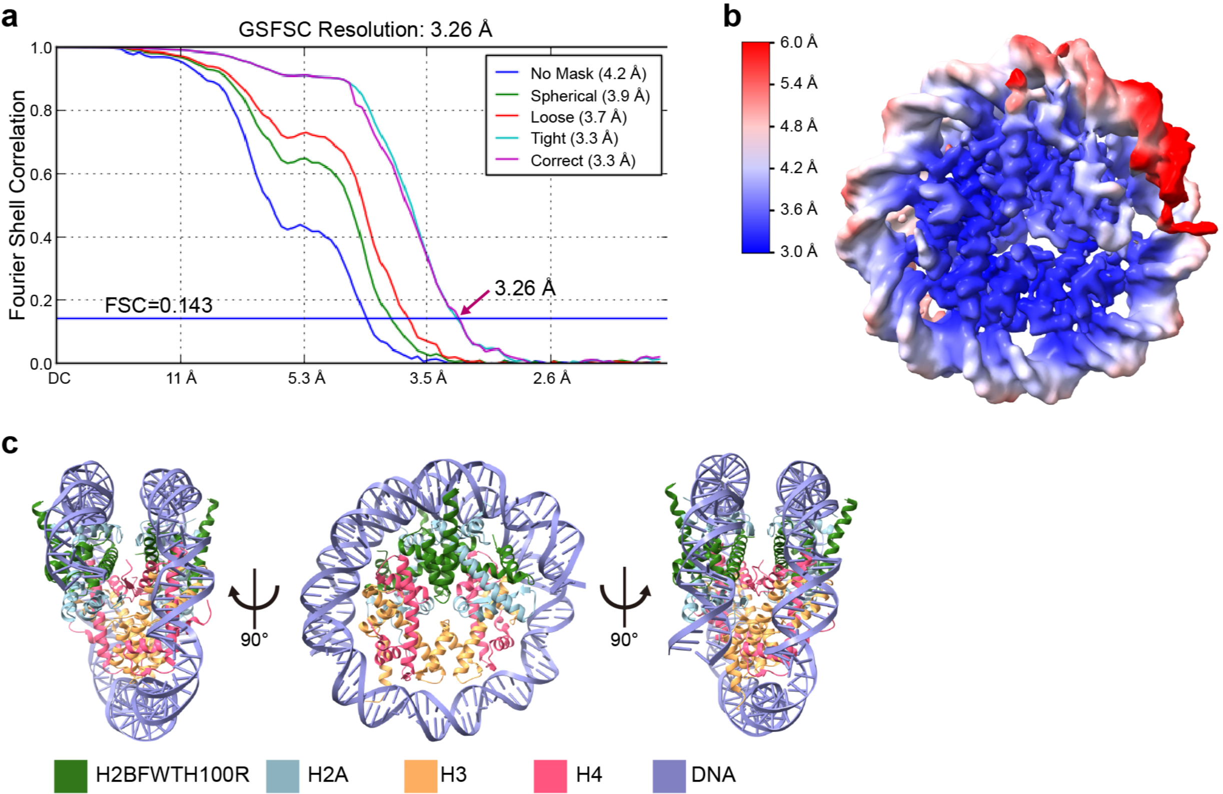

Figure S10 Ding *et al.*,

a

b

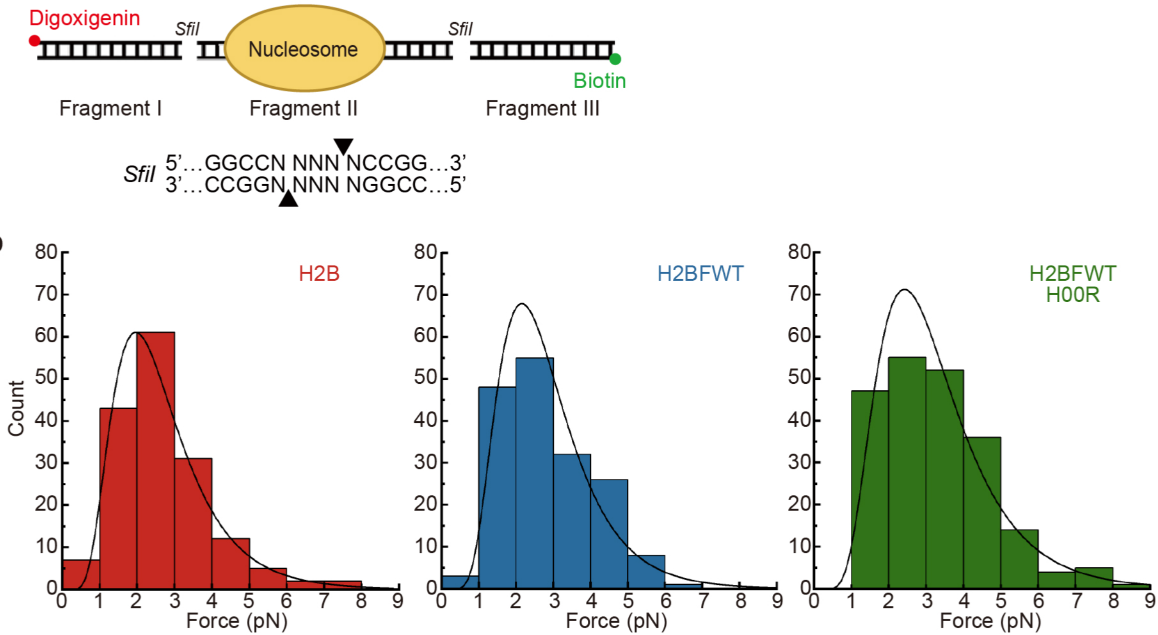


c

| | Inner Unwrap | | Inner Rewrap | |
|---|---|---|---|---|
| | Force (pN) | n | Force (pN) | n |
| H2B | 14.4±0.14 | 434 | 2.7±0.11 | 163 |
| H2BFWT | 10.8±0.14 | 236 | 2.83±0.09 | 173 |
| H2BFWT H100R | 11.1±0.11 | 311 | 3.25±0.10 | 214 |

d

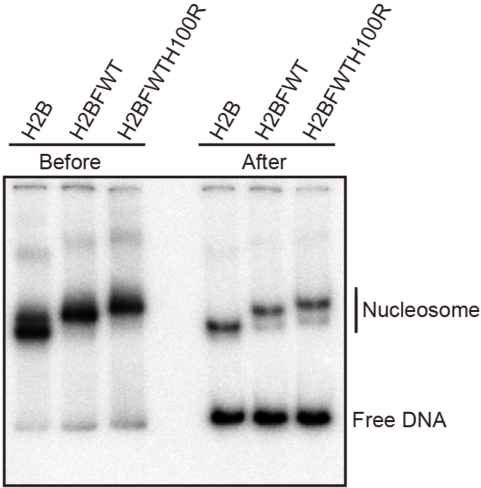

### Figure S11 Ding *et al.*,

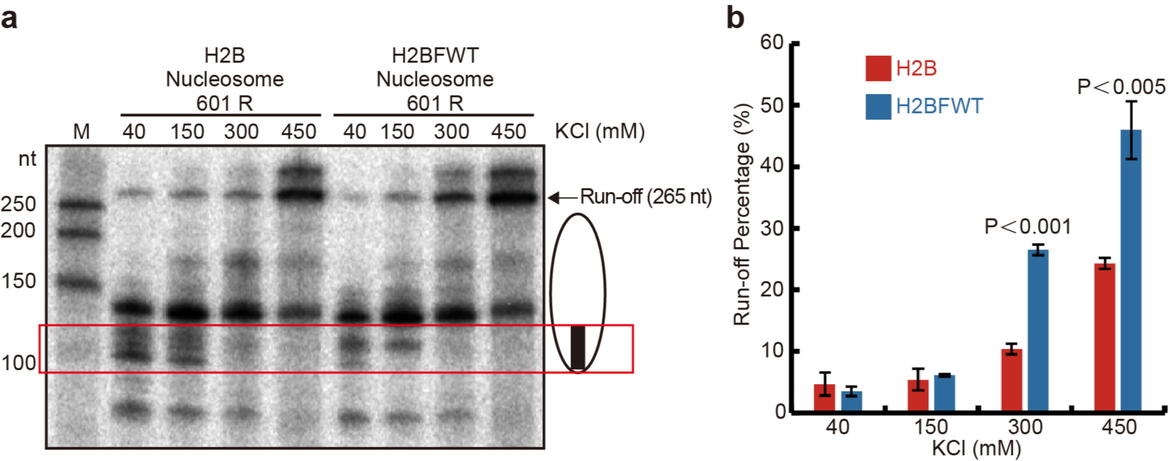

**Table S1. List of hydrogen-bond-forming amino acid residues between H2A-H2B/H2BFWT dimer and DNA**

| | H2B Nucleosome | | H2BFWT Nucleosome | |
|---|---|---|---|---|
| | Atom A | Atom B | Atom A | Atom B |
| 1 | H2A R17 N | I DT -43 OP1 | H2A R17 N | I DA -43 OP1 |
| 2 | H2A R32 NE | I DA -44 OP1 | H2A V43 N | J DA 39 OP1 |
| 3 | H2A V43 N | J DA 39 OP1 | H2A A45 N | J DT 38 OP1 |
| 4 | H2A A45 N | J DG 38 OP1 | H2A T76 N | J DG 58 OP1 |
| 5 | H2A K75 NZ | J DA 59 OP2 | H2A T76 OG1 | J DG 58 OP1 |
| 6 | H2A T76 N | J DC 58 OP1 | H2BFWT R77 NH1 | I DC -54 OP2 |
| 7 | H2A T76 OG1 | J DG 57 O3’ | H2BFWT Q108 N | I DA -34 OP1 |
| 8 | H2B Y40 OH | J DG 48 OP1 | H2BFWT T109 OG1 | I DA -34 OP1 |
| 9 | H2B Y40 OH | J DG 48 O5’ | H2A’ R20 NH2 | J DT -42 OP2 |
| 10 | H2B Y42 OH | I DG -53 O5’ | H2A’ R29 NH2 | I DC 49 OP1 |
| 11 | H2B S56 N | I DA -54 OP1 | H2A’ R32 NE | J DA -44 OP1 |
| 12 | H2B S87 N | I DG -34 OP1 | H2A’ R42 NH2 | I DG 38 O4’ |
| 13 | H2A’ K13 NZ | J DA -41 OP1 | H2A’ V43 N | I DA 39 OP1 |
| 14 | H2A’ K13 NZ | J DG -42 O3’ | H2A’ T76 N | I DC 58 OP1 |
| 15 | H2A’ R17 N | J DA -43 OP1 | H2BFWT’ Y61 OH | I DG 48 OP1 |
| 16 | H2A’ R20 NH2 | J DG -42 OP1 | H2BFWT’ R77 N | J DA -54 OP1 |
| 17 | H2A’ R29 NH2 | I DC 49 OP1 | H2BFWT’ Q108 N | J DG -34 OP1 |
| 18 | H2A’ V43 N | I DA 39 OP1 | H2BFWT’ T109 N | J DG -34 OP1 |
| 16 | H2A’ A45 N | I DT 38 OP1 | | |
| 20 | H2A’ T76 N | I DG 58 OP1 | | |
| 21 | H2B’ R33 N | I DA 50 OP1 | | |
| 22 | H2B’ Y40 OH | I DG 48 OP1 | | |
| 23 | H2B’ Y42 OH | J DA -53 O5’ | | |
| 24 | H2B’ I54 N | J DA -53 OP1 | | |
| 25 | H2B’ S56 N | J DC -54 OP1 | | |
| 26 | H2B’ S87 N | J DA -34 OP1 | | |
| 27 | H2B’ S87 OG | J DA -34 OP1 | | |
| Total | 27 | | 18 | |

**Table S2 List of hydrogen-bond-forming amino acid residues betweenH2A-H2B/H2BFWT dimer and H3-H4 tetramer**

| | H2B Nucleosome | | H2BFWT Nucleosome | |
|---|---|---|---|---|
| | Atom A | Atom B | Atom A | Atom B |
| 1 | H2A T101 N | H4’ T96 O | H2A R99 NH1 | H4’ G94 O |
| 2 | H2A N110 N | H3’ Q55 OE1 | H2BFWT W113 NE1 | H4 H75 O |
| 3 | H2A N110 O | H3’ Q55 NE2 | H2BFWT E114 OE2 | H4 H75 NE2 |
| 4 | H2A N110 OD1 | H3’ Q55 NE2 | H2A’ R81 NH2 | H3 K56 O |
| 5 | H2B R92 NH2 | H4 E74 O | H2A’ R99 O | H4 T96 N |
| 6 | H2B R92 NH1 | H4 H75 O | H2A’ T101 N | H4 T96 O |
| 7 | H2A’ T101 N | H4 T96 O | H2A’ Q104 N | H3 E94 OE1 |
| 8 | H2A’ T101 O | H4 Y98 N | H2A’ N110 N | H3 Q55 OE1 |
| 9 | H2A’ N110 N | H3 Q55 OE1 | H2BFWT’ W113 NE1 | H4’ H75 O |
| 10 | H2A’ L115 | H4 K44 NZ | | |
| 11 | H2B’ D68 OD2 | H4 Y98 OH | | |
| 12 | H2B’ R92 NH2 | H4’ E74 O | | |
| 13 | H2B’ R92 NE | H4’ H75 O | | |
| 14 | H2B’ E93 OE1 | H4’ H75 NE2 | | |
| 15 | H2B’ T96 OG1 | H4’ H75 ND1 | | |
| Total | 15 | | 9 | |

**Table S3. List of hydrogen-bond-forming amino acid residues betweenH3-H4 tetramer and DNA**

| | H2B Nucleosome | | H2BFWT Nucleosome | |
|---|---|---|---|---|
| | Atom A | Atom B | Atom A | Atom B |
| 1 | H3 R40 NH2 | J DT 9 O2 | H3 R42 NH1 | I DG -6 O3’ |
| 2 | H3 R42 NH2 | I DA -5 OP1 | H3 L65 N | J DC 18 OP2 |
| 3 | H3 G44 N | J DT 9 OP1 | H3 R69 NH2 | J DA 17 OP2 |
| 4 | H3 A47 N | J DT 9 OP1 | H3 R72 NH1 | I DT -23 OP1 |
| 5 | H3 R49 NE | J DT -65 OP1 | H3 R72 NH2 | I DT -23 OP1 |
| 6 | H3 R63 NH2 | J DG 18 OP1 | H3 R83 NH2 | I DT -24 O3’ |
| 7 | H3 R63 NE | J DG 18 OP1 | H3 F84 N | I DT -23 OP1 |
| 8 | H3 R63 NE | J DA 17 O3’ | H3 V117 N | I DA -3 OP1 |
| 9 | H3 K64 N | J DG 18 OP1 | H3 T118 N | I DA -3 OP1 |
| 10 | H3 L65 N | J DG 18 OP2 | H3 T118 OG1 | I DA -3 OP1 |
| 11 | H3 R72 NH1 | I DC -23 OP1 | H4 I46 N | J DC 8 OP1 |
| 12 | H3 R72 NH2 | I DC -23 OP2 | H4 K79 N | J DG 28 OP1 |
| 13 | H3 F84 N | I DC -23 OP1 | H4 T80 OG1 | J DG 28 OP1 |
| 14 | H4 R35 NH1 | J DG 8 OP2 | H3’ R63 NH1 | J DA -13 OP1 |
| 15 | H4 R35 NH2 | J DG 8 OP2 | H3’ R63 NH1 | J DA -14 O3’ |
| 16 | H4 R36 NH1 | I DA -13 OP1 | H3’ R63 NH2 | J DA -13 OP1 |
| 17 | H4 R36 NH2 | I DA -13 OP1 | H3’ R63 NH2 | J DA -14 O3’ |
| 18 | H4 I46 N | J DG 8 OP1 | H3’ K64 N | I DG 18 OP1 |
| 19 | H4 G48 N | J DC 7 OP1 | H3’ L65 N | I DG 18 OP2 |
| 20 | H3’ R42 N | J DC 70 OP1 | H3’ R72 NH1 | J DC -23 OP1 |
| 21 | H3’ R42 NH1 | J DG -5 OP1 | H3’ R72 NH2 | J DC -23 OP2 |
| 22 | H3’ G44 N | I DG 9 OP1 | H3’ F84 N | J DC -23 OP1 |
| 23 | H3’ T45 OG1 | J DC 70 OP1 | H3’ S86 N | J DG -24 OP2 |
| 24 | H3’ A47 N | I DG 9 OP1 | H3’ T118 OG1 | J DG -3 OP1 |
| 25 | H3’ K64 N | I DC 18 OP1 | H4’ T30 OG1 | J DC -12 OP1 |
| 26 | H3’ L65 N | I DC 18 OP2 | H4’ R35 NH1 | I DG 8 OP2 |
| 27 | H3’ R69 NH1 | I DA 17 OP2 | H4’ I46 N | I DG 8 OP1 |
| 28 | H3’ R69 NH2 | I DA 17 OP2 | H4’ G48 N | I DC 7 OP1 |
| 29 | H3’ R72 NH1 | J DT -23 OP1 | H4’ T80 N | I DA 28 OP1 |
| 30 | H3’ R72 NH2 | J DT -23 OP2 | | |
| 31 | H3’ F84 N | J DT -23 OP1 | | |
| 32 | H3’ V117 N | J DA -3 OP1 | | |
| 33 | H3’ T118 N | J DA -3 OP1 | | |
| 34 | H4’ T30 OG1 | J DA -12 OP2 | | |
| 35 | H4’ R36 NH1 | J DA -13 OP2 | | |
| 36 | H4’ R36 NH2 | J DA -13 OP1 | | |
| 37 | H4’ I46 N | I DC 8 OP1 | | |
| 38 | H4’ K79 NZ | I DG 27 OP1 | | |
| 39 | H4’ T80 OG1 | I DG 28 OP1 | | |
| 40 | H4’ T80 OG1 | I DG 28 O5’ | | |
| Total | 40 | | 29 | |

**Table S4. Oligos or DNA templates used in this study**

| Name | Sequence (5'-3') |
|---|---|
| GAPDH-F (ChIP-qPCR) | TCGACAGTCAGCCGCATCT |
| GAPDH-R (ChIP-qPCR) | CTAGCCTCCCGGGTTTCTCT |
| MYOD1-F | CCGCCTGAGCAAAGTAAATGA |
| MYOD1-R | GGCAACCGCTGGTTTGG |
| SAT2-F | ATCGAATGGAAATGAAAGGAGTCA |
| SAT2-R | GACCATTCGATGATTGCATTCA |
| H2A.P-F | CGTCCAAAGCCAGAGTTCCT |
| H2A.P-R | TGCGCATACTGCCATTGTTG |
| TH2B-F | CCGGAGGTGTCATCTAAAGGT |
| TH2B-R | CTCCTTACGGGTCCTCTTGC |
| HOXA11-F | CTAGGTGCCCAGAATGAGGC |
| HOXA11-R | AGGATTTGCAGCCCTCTTCC |
| GAPDH-F (RT-qPCR) | GAAGGTGAAGGTCGGAGTCAAC |
| GAPDH-R (RT-qPCR) | CAGAGTTAAAAGCAGCCCTGGT |
| HAUS3-F | GGTTGTTTGAGGGCGTTGAA |
| HAUS3-R | TCCAATGCCGCCCCTTCTA |
| POLN-F1 | CAATGGACCTTTGCTCTAAACTG |
| POLN-R1 | CCGTTCTCCTGCAACAAAAT |
| POLN-F2 | TACAACTTGAAGACAGGAAGACTC |
| POLN-R2 | GATCTGTGAGCCTGACACTGAAG |
| POLN-F3 | AATCAAGGACTTCGCCCGAG |
| POLN-R3 | CCTTGCACCACGAAGTTCAC |
| Fragment 1 Forward | Digoxin-GGTGATGACGGTGAAAACCT |
| Fragment 1 Reverse | CGGCCACATTGGCCTTGCCGGTGCCAGTCGGATA |
| Fragment 2 Forward | CGGCCAATGTGGCCGGTCGCTGTTCAATACATGC |
| Fragment 2 Reverse | CGGCCTACGAGGCCTGTGATGGACCCTATACG |
| Fragment 3 Forward | CGGCCTCGTAGGCCAAGCTAGAGTAAGTAGTT |
| Fragment 3 Reverse | Biotin-ACGAAAGGGCCTCGTGATAC |
| TDS | CTCTTCCCGGGTTCGCCTTGTCTCGGGCGTCGGCTGTAAGTATCCTATACC |
| NDS | CCGACGGTATAGGATACTTACAGCCGACGCCCGAGACAAGGCGAACCCGGGAAGAGCAT |
| RNA14 | UUUUUGACGCCCGA |
| 601.2* | GGACCCTATACGC**GGCC**GCCCTGCAGAAGCTTGGTCCCGG**GGCC**GCTCAATTGGTCGTAGCAAGCTCTGGATCCGCTTGATCGAACGTACGCGCTGTCCCCCGCGTTTTAACCGCCAAGGGGATTACTCCCTAGTCTCCAGGCACGTGTCAGATATATACATCCTGTGCATGTA |
| 601.2 *Hae*III* | GGACCCTATACGCGGCGCCC**GGCC**AGAAGC**GGCC**TCCCGG**GGCC**GCTCAA**GGCC**TCGTAG**GGCC**CTCTGG**GGCC**GCTTGA**GGCC**ACGTACG**GGCC**GTCCCCCGCGTTTTAACCGCCAAGGGGATTACTCCCTAGTCTCCAGGCACGTGTCAGATATATACATCCTGTGCATGTA |

* *Hae*III restriction digestion sites are labeled with bold.
