## Supplemental Information for "Testis-specific H2BFWT disrupts nucleosome integrity through reductions of DNA-histone interactions"

### Supplemental material

### H2BFWT antibody quality test

An anti-H2BFWT antibody was raised in rabbits using the H2BFWT peptide (36-49 aa, SKQRKRGRHGPRRC). To check the quality of the antibody, the ChIP qPCR experiment was performed using stable Flag-tagged H2BFWT HeLa cell lines.

Nuclear extracts prepared from the stable Flag-H2BFWT HeLa cells induced with different amounts of doxycycline were loaded onto a 15% SDS-PAGE and Western blotting was performed to check the antibody specificity. The ChIP experiment was performed based on Pierce™ Agarose ChIP Kit (Thermo Fisher Scientific) following the manufacture's instruction. Flag monoclonal antibody (Sigma-Aldrich) was used as the positive control, and the homemade anti-H2BFWT antibody was used to perform the ChIP assay. ChIP DNA was amplified with gene-specific primers and quantified based on LightCycler® 480 SYBR Green I Master (Roche) with different primers (Table S4).

### Single-cell sequencing data analysis

Human testis single-cell RNA sequencing data was kindly provided from Prof. Jie Qiao (Peking University). The single cell transcriptome TPM matrix was imported into seurat R package to start the analysis pipeline. The expression data was normalized by dividing the TPM values with 100 and applying log1p to the results of the divisions. The variable genes were identified by the following parameters: x.low.cutoff of 0.05, x.high.cutoff of 3, and y.cutoff of 0.5. In the dimension reduction analysis, the first 20 PCs (principle components) were subjected into the downstream UMAP (Uniform Manifold Approximation and Projection) analysis. Next, we selected the first 10 PCs and resolution of 0.6 to implement the cell clustering analysis. Cluster markers were identified by Findmarkers function. We then manually projected the calculated cell clusters to the real cell types based on marker genes expression. Gene expression visualization was carried out by the Featureplot function of the Seurat package.

### Chromatin immunoprecipitation from human testis and sequencing library preparation

The ChIP assay from human testis was followed by the published methods with modifications [1,2]. FFPE normal tissue blocks of human testis were purchased from Origene (CB811079). Testis blocks were sliced into 10-μm thickness sections. Paraffin was removed and tissue sections were rehydrated with a series of xylene and ethanol/water solution. Twenty testis tissue sections were removed from the slides and kept in a resuspension buffer (50 mM Tris-HCl (pH 7.5), 150 mM NaCl, 0.5% Tween-20, 20 ng/μL RNase A) for 30 min at RT. After centrifugation at 18,000 g for 5 mins, the pellet was resuspended in 1 mL of a buffer (50 mM Tris-HCl (pH 7.5), 4 mM $MgCl_2$, 1 mM $CaCl_2$) with 10 units/mL MNase (Worthington) and the incubated at

37°C for 9 min and then the reaction was stopped by adding 15 mM of EDTA. Thermolabile Proteinase K (NEB, P8111S) was added to 0.08 unit/mL at 37°C for 6 min after incubation of the samples at 40°C for 1 h. The reaction was inactivated at 65°C for 15 min. After further fragmentation with sonication, the samples were centrifuged at 21,000 g for 5 min at 4°C. Three volumes of Ch-IP buffer (30 mM Tris-HCl (pH 7.5), 50 mM NaCl, 3 mM EDTA, and 0.1 mM PMSF) was added to the supernatant. Antibody against H2BFWT (25 μg), which was raised from the synthesized peptide (amino acids, 36-49, SKQRKRGRHGPRRC, Shanghai Youke Biotechnology) was added to the supernatant and incubated for 16 h at 4°C with rotation. 15 μL ChIP-grade protein A/G Plus Agarose (Thermo Scientific, 26159) was blocked by BSA for 16 h at 4°C.

The beads and antibody-incubated sample were further incubated for 3 h at 4°C. After centrifugation at 2,000 g for 2 min at 4°C, the pellet was washed sequentially with 1 mL washing buffer A/B/C (50/100/150 mM NaCl, 50 mM Tris-HCl (pH 7.5), 1% TritonX-100, 5 mM EDTA). TE buffer containing 200 mM NaCl was added to the beads with a further incubation at 65°C for 20 hrs for de-cross-linking. Proteinase K (NEB, P8107S) was added to 80 ng/μL with incubation at 45°C for 3 h. The DNA was extracted, and DNA libraries were prepared with Smarter Thruplex DNAseq Kit (Clontech, R400674) following the manufacturer's protocols with amplification for 10-13 circles. Sequencing was performed based on Nextseq 500 instrument (Illumina) with paired-end 150 bp (PE150) sequencing method.

**Chromatin immunoprecipitation sequencing result analysis**

The quality of the clean read was checked with FastQC (Galaxy Version 0.72+galaxy1). Then, the reads were trimmed to remove the adapter sequence with Trimmomatic (Galaxy Version 0.38.0). The trimmed reads were mapped onto the human genome (hg19) with Bowtie 2 (Galaxy Version 2.3.4.3+galaxy0). The filtered input and H2BFWT bam files were converted to be bed files. Random DNA sequence and mitochondrial DNA were removed with Select (Galaxy Version 1.0.1). The peaking calling was performed with MACS2 callpeak (Galaxy Version 2.1.1.20160309.6). To obtain the gene list of the H2BFWT enriched gene list, the GREAT (version 4.0.4) was used to make the prediction. H2BFWT genome wide distribution and genomic annotation were plotted with ChIPSeeker [3]. Heatmap of the peaks and heatmap around the TSS, TES and gene body was plotted with EaSeq [4]. The peaks shown in the genome are based on IGB (Version 9.1.6).

**Quantitative reverse transcription PCR analysis (RT-qPCR) of stable Flag-H2BFWT HeLa cells**

Stable Flag-H2BFWT HeLa cells were induced with the indicated concentration of doxycycline at 37ºC and 5% $CO_2$ for 48 hours. Total RNAs were extracted using the PureLink™ RNA Mini Kit (Invitrogen) and cDNA libraries were subsequently synthesized using the SuperScript™ IV First-Strand Synthesis System (Invitrogen) with oligo $dT_{20}$ primer according to the manufacturer's recommended protocols. Expressions of *POLN* and *HAUS3* were detected using gene-specific primers with LightCycler® 480 SYBR Green I Master (Roche) on a LightCycler-® 480 Instrument II (Roche). The annealing temperature of the real-time PCR reaction was set at 60ºC. The relative expression of each target gene was calculated using the delta-delta-Ct method with the Ct value of GAPDH (RT-qPCR) primers as the internal housekeeping gene control. The cDNA library of un-induced cells was used as the baseline control.

**Histone expression, purification, and nucleosome loading**

H2BFWT recombinant protein purification followed the same method as canonical histone purification with modification [5]. His-H2BFWT was purified using Ni-NTA resin (Qiagen) under urea denaturing condition. The His-tag was removed by thrombin digestion. After the digestion, Ni-NTA resin was used to remove the His-tag and undigested proteins. The sample was further purified by C-18 reverse-phase chromatography (Agilent) with acetonitrile buffer containing 0.1% trifluoroacetic acid.

To load nucleosomes, 601 nucleosome positioning DNA sequence was used [6]. For the optical tweezers single-nucleosome assay, three DNA fragments were used, and they can be ligated (Figure S10A). All the three DNA fragments were amplified by PCR and digested by *Sfi*I (NEB). PCR products and digested DNA were purified based on ethanol and PEG 8000 precipitation to reduce the damage on the DNA.

**Nucleosome salt stability assay**

Amicon Ultra Centrifugal Filters (50 kDa cut-off) (Millipore) was used to concentrate the purified nucleosomes to 200 ng/μL. Nucleosome (2.5 μL) and different salt concentration buffers (2.5 μL) (final NaCl concentrations were indicated in the Figure S6; 10 mM Tris, pH 7.5, 0.5 mM EDTA, and 5 mM 2-Mercaptoethanol) were mixed. After incubation on ice for 10 min, the samples were dialyzed against 0 M nucleosome loading buffer for 5 min on the surface of a 47-mm MF-Millipore membrane (Millipore, VSWP04700) to remove the salt. The products were checked with 5% Native-PAGE, 1X TBE.

**Pol II Transcription elongation assay with nucleosomal DNA template**

*S. cerevisiae* Pol II containing a hexahistidine-tagged Rpb3 subunit (strain kindly provided by the laboratory of Patrick Cramer) was purified as described [7]. 4 pmol of purified *S. cerevisiae* Pol II was immobilized on 50 μL Nickel-NTA beads (Qiagen). RNA and template DNA (TDS) scaffold was assembled by annealing the template strand DNA (25 pmol) and the [$\gamma$-$^{32}$P]-ATP end-labeled 14 mer RNA (RNA14, 36 pmol) by linear gradient from 42 °C to room temperature [8]. Transcription elongation complex (TEC) was formed by incubating the scaffold with Pol II coupled agarose at room temperature for 10 min followed by the addition of non-template strand (NDS, 37.5 pmol) at 37 °C for 10 min. The TEC was washed with TB40 (20 mM Tris-HCl (pH 7.9), 5 mM $MgCl_2$, 5 mM 2-Mercaptoethanol, 10 μM $ZnCl_2$, and 40 mM KCl) and ligated to the 601R (601 Reverse DNA template) nucleosome through *Dra*III site at 16 °C for 1 hr. Then the complex was transcribed in the presence of the indicated concentrations of KCl and 500 µM rNTPs. Transcription reactions were stopped by adding 20 µg proteinase K and incubated at 30°C for 20 min. Urea stop buffer (89 mM Tris-HCl (pH 8.0), 8 M urea, 89 mM boric acid, 50 mM EDTA, 0.25% bromophenol blue) was further added and the reactions were incubated at 65°C for 20 min. Transcripts were resolved on 12% denaturing polyacrylamide gels. Gels were scanned with Sapphire Biomolecular Imager (Azure Biosystem).

Mammalian Pol II was purified as described previously [9]. Briefly, 1 pmol of purified mammalian Pol II was immobilized on 8WG16-Agarose Protein A beads. The following procedure was the same as in the yeast Pol II transcription assay with the decreasing the TDS to 10 pmol, the RNA14 to 11.25 pmol and the NDS to 15 pmol. Gels were scanned and quantified by a Typhoon phosphorimager (GE healthcare).

**Extraction of $\Delta$□$^{0}$ from the nucleosome hopping data.**

The free energy $\Delta G^0$ of the nucleosome outer DNA wrap was calculated based on the following formula [10-12]:

$$\Delta G^0 = F_{eq}\Delta x - \Delta G_{strech} - k_B T \ln k_{eq}$$

The equilibrium force ($F_{eq}$) is the force at which the wrapping rate ($k_w$) equals the unwrapping rate ($k_u$). The $k_w$ or $k_u$ is the reciprocal of the keeping time (second) of a given unwrapped state or wrapped state. The $\Delta x$ is the average extension change of the outer unwrap in the nucleosome pulling assay. $\Delta G_{stretch}$ is the energy needed to stretch the free DNA template to the $F_{eq}$. The equilibrium rate ($k_{eq}$) is $k_w$ or $k_u$ at $F_{eq}$. $k_B T$ is the product of the Boltzmann constant ($k_B$) and the temperature ($T$).

### Supplemental Figure legends

#### Figure S1 Amino acid sequence alignment and single-cell RNA expression analysis

(A) Amino Acid sequence alignment of H2B, TH2B and H2BFWT. Underline and wavy line indicate the peptide epitopes for TH2B and H2BFWT antibodies generation, respectively. Redbox indicates the position of the H2BFWTH100R SNP.

(B) Single-cell RNA expression patterns of spermatogenesis marker genes, H2BFWT and TH2B during spermatogenesis.

#### Figure S2 ChIP-qPCR validation of the H2BFWT antibody

(A) Western blot detecting the Flag-H2BFWT expression in Flag-H2BFW stable HeLa cell nuclear extract induced with different concentrations of doxycycline. WT

HeLa cell nuclear extraction and 0 μg/mL doxycycline were used as negative controls.

(B) The procedure of ChIP-experiment using Flag-H2BFWT stable HeLa cells.

(C) ChIP DNA recovery percentage from stable Flag-H2BFWT HeLa cell. qPCR was performed to detect the DNA recovery percentage (% input) under different doxycycline induction conditions. Anti-Flag antibody was used as the positive control, and both no antibody and 0 ng/mL doxycycline conditions were used as negative controls. Error bar indicates SD (n=3).

**Figure S3 The genome wide profiling of H2BFWT peaks in the human chromosome**

**Figure S4 Heatmaps of H2BFWT enrichment region alignments**

(A) Heatmap of H2BFWT binding sites within 10 kb of H2BFWT peaks. After MACS2 peak calling, the H2BFWT enrichment regions ($p<0.05$) were plotted within ±5 kb of the peaks.

(B) Heatmap of H2BFWT enrichment around the transcription end sites (TES) (±5 kb) of all the genes.

(C) Heatmap of H2BFWT binding sites within the gene body (±100%, relatively).

**Figure S5 *HAUS3* and *POLN* expressions are induced by the presence of H2BFWT**

(A) Heat map illustration of genes highly expressed in testis of H2BFWT enriched the highest 200 peaks.

(B) The expression patterns of *HAUS3* and *POLN* in single-cell RNA sequence analysis during spermatogenesis.

(C) ChIP-qPCR of *HAUS3* and *POLN* in the presence or absence of H2BFWT. The expression level was adjusted according to the GAPDH q-PCR signal.

**Figure S6 Salt stabilities of H2B, H2BFWT and H2BFWH100R nucleosomes**

(A) Representative gel of the salt stability assay of H2BFWT and H2B nucleosomes. Stars represent nucleosome-nucleosomes and histone-DNA complexes.

(B) Relative free DNA percentage in each lane of (A). The error bars represent SEM (n=5).

(C) Representative gel of the salt stability assay of H2BFWT and H2BFWTH100R nucleosomes. Stars represent the nucleosome-nucleosomes and histone-DNA complexes.

(D) Relative free DNA percentage in each lane of (C). The error bars represent SEM (n=5).

**Figure S7 One-pot assay analysis in time course**

(A-C) Relative digestion efficiencies of H2BFWT and H2BFWTH100R nucleosomes against H2B nucleosomed were calculated for samples incubated with *Hae*III for (A) 10 mins, (B) 20 mins, and (C) 40 mins.

(D-F) Relative digestion efficiencies of H2BFWTH100R nucleosomes against H2BFWT nucleosomes were calculated for samples incubated with *Hae*III for (D) 10 mins, (E) 20 mins, and (F) 40 mins.

**Figure S8 Cryo-EM data processing**

(A) Representative cryo-EM micrograph (left) and 2D class averages (right) of H2BFWT nucleosome particles.
(B) Flowchart of cryo-EM data processing for the H2BFWT nucleosome.
(C) Fourier Shell Correlation curve of the H2BFWT nucleosome density map. The final resolution is 3.49 Å (purple).
(D) The final 3D density map of the H2BFWT nucleosome with the local resolution overlay.

**Figure S9 H2BFWTH100R nucleosome structure model and its resolution distribution map**

(A) Fourier Shell Correlation curve of the H2BFWTH100R nucleosome density map. The final resolution is 3.26 A (purple).
(B) The final 3D density map of the H2BFWTH100R nucleosome with the local resolution overlay.
(C) Overall structures of the H2BFWTH100R nucleosome atomic model in disc view (middle), and gyre view (left and right).

**Figure S10 Inner rip force distribution and nucleosome stability property of in the single-molecule optical tweezers assay**

(A) Geometry of the tether-DNA construct. Fragment II contains a 601 nucleosome positioning sequence.
(B) Histograms of inner rewrap force among H2B, H2BFWT, H2BFWTH100R nucleosomes. The black lines represent the best-fitting regression curves.
(C) The summary of inner unwrap and rewrap mean forces statistics. The errors indicate ±1 SEM.
(D) H2B, H2BFWT and H2BFWTH100R nucleosomes with $^{32}$P-labeled DNA were analyzed before and after incubation at 1 ng/μL concentration in 300 mM NaCl for 10 mins by 5% native PAGE.

**Figure S11 Mammalian Pol II transcription elongation assay**

(A) Nucleosomes-containing templates were transcribed by mammalian Pol II in the presence of the indicated concentrations of KCl. Arrow indicates the position of Run-off transcripts. The position of the nucleosome on the template is indicated by the oval (the entry or H2A/H2B dimer region is shown as a black square and highlighted with a red frame).
(B) The percentage of run-off transcripts was quantified under different salt conditions. Error bars represent ±1 SEM in repeated experiments (n=4).

**Table S1 List of hydrogen-bond-forming amino acid residues between H2A-H2B/H2BFWT dimer and DNA**

**Table S2 List of hydrogen-bond-forming amino acid residues betweenH2A-H2B/H2BFWT dimer and H3-H4 tetramer**

**Table S3 List of hydrogen-bond-forming amino acid residues betweenH3-H4 tetramer and DNA**

**Table S4 The primers and DNA sequence that used for this study**

**References**


1. Cejas, P. et al. Chromatin immunoprecipitation from fixed clinical tissues reveals tumor-specific enhancer profiles. *Nature medicine* **22**, 685-691 (2016).
2. Fanelli, M. et al. Pathology tissue–chromatin immunoprecipitation, coupled with high-throughput sequencing, allows the epigenetic profiling of patient samples. *Proceedings of the National Academy of Sciences* **107**, 21535-21540 (2010).
3. Yu, G., Wang, L.-G. & He, Q.-Y. ChIPseeker: an R/Bioconductor package for ChIP peak annotation, comparison and visualization. *Bioinformatics* **31**, 2382-2383 (2015).
4. Lerdrup, M., Johansen, J.V., Agrawal-Singh, S. & Hansen, K. An interactive environment for agile analysis and visualization of ChIP-sequencing data. *Nature structural & molecular biology* **23**, 349-357 (2016).
5. Wittmeyer, J., Saha, A. & Cairns, B. DNA translocation and nucleosome remodeling assays by the RSC chromatin remodeling complex. *Methods in enzymology* **377**, 322-343 (2004).
6. Lowary, P. & Widom, J. New DNA sequence rules for high affinity binding to histone octamer and sequence-directed nucleosome positioning. *Journal of molecular biology* **276**, 19-42 (1998).
7. Sydow, J.F. et al. Structural basis of transcription: mismatch-specific fidelity mechanisms and paused RNA polymerase II with frayed RNA. *Molecular cell* **34**, 710-721 (2009).
8. Kireeva, M.L., Lubkowska, L., Komissarova, N. & Kashlev, M. Assays and affinity purification of biotinylated and nonbiotinylated forms of double-tagged core RNA polymerase II from Saccharomyces cerevisiae. *Methods in enzymology* **370**, 138-155 (2003).
9. Wan, Y.C.E. et al. Cancer-associated histone mutation H2BG53D disrupts DNA–histone octamer interaction and promotes oncogenic phenotypes. *Signal transduction and targeted therapy* **5**, 1-4 (2020).
10. Bustamante, C., Chemla, Y.R., Forde, N.R. & Izhaky, D. Mechanical processes in biochemistry. *Annual review of biochemistry* **73**, 705-748 (2004).
11. Liphardt, J., Onoa, B., Smith, S.B., Tinoco Jr, I. & Bustamante, C. Reversible unfolding of single RNA molecules by mechanical force. *Science* **292**, 733-737 (2001).
12. Li, W. et al. FACT remodels the tetranucleosomal unit of chromatin fibers for gene transcription. *Molecular cell* **64**, 120-133 (2016).
